# The evolution of tandem repeat sequences under partial selfing and different modes of selection

**DOI:** 10.1101/2025.07.04.663195

**Authors:** Vitor Sudbrack, Charles Mullon

## Abstract

Tandem repeat sequences (TRs) occur when short DNA motifs are repeated head-to-tail along chromosomes and are a major source of genetic variation. Population genetics models of TR evolution have focused on large, randomly mating, haploid populations. Yet many organisms reproduce partially through self-fertilisation (“selfing”), which increases homozygosity and thus may alter the evolutionary processes shaping TRs. Here we use mathematical modelling and simulations to study the evolution of homologous TRs in partially selfing, diploid populations under four different selective regimes that may be relevant to TRs: (i) additive purifying selection, (ii) truncation-like purifying selection, (iii) selection against heterozygotes due to misalignment costs, and (iv) stabilising selection favouring an intermediate TR length. We show that selfing influences TR evolution primarily by increasing homozygosity, with two main consequences: (1) it enhances the variation produced by unequal recombination within individuals, and (2) it increases variation between individuals. Consequently, selection against TR expansions becomes more effective under partial selfing across all modes of selection considered, resulting in shorter TRs and lower genetic load, despite higher genetic drift. Overall, our results suggest that mating systems and inbreeding are important factors shaping variation in TRs.

## 1 Introduction

Tandem repeat sequences (TRs) consist in short genomic motifs, varying between 2 to 2’000 base pairs, that are repeated head-to-tail from a dozen to thousands of times (e.g. GCCGCCGCC…), spanning up to millions of base pairs in some cases (Miklos and Gill, 1982; Depienne and Mandel, 2021). They are ubiquitous across the tree of life (Verbiest et al., 2023), including microsatellite (1-8 base pair motifs), minisatellite (9-99 base pairs motifs), and satellite DNA (>100 base pairs motifs, Jeffreys et al., 1985; Buschiazzo and Gemmell, 2006; Richard et al., 2008). The number of repeated motifs within a TRs, called “TR copy number” or “TRs length”, varies greatly among populations and among individuals of the same population, making them useful genetic markers (in e.g. population genetics, paternity testing, and linkage mapping; Hammond et al., 1994; Slatkin, 1995; Verbiest et al., 2023). For long, TRs have been considered as part of “junk DNA”, moulded either by mutational processes only or in combination with purifying selection (Charlesworth et al., 1994; Kruglyak et al., 1998, 2000; McGinty et al., 2025). Recent studies however have revealed that at least some of these sequences have functional significance and may therefore play an adaptive role (John and Miklos, 1979; Kruglyak et al., 2000; Balzano et al., 2021; Depienne and Mandel, 2021; Verbiest et al., 2023). Understanding the processes that shape variation in TRs is therefore useful for both practical and fundamental reasons.

Population genetic models have been useful to disentangle the effects of selection, mutation, recombination and genetic drift on the evolution of TRs (Ohta, 1983b; Stephan, 1986, 1987; Walsh, 1987; Stephan, 1989; see also Crow and Kimura, 1970 p. 294-296; Krüger and Vogel, 1975; Takahata, 1981; Ohta, 1983a for models of chromosome size or gene content evolution that involve similar processes). This body of work has highlighted how the particular mechanisms of mutation and recombination affecting TRs contribute to variation in TRs length. Specifically, the repeated structure of these sequences makes them subject to replication slippage (a mutational process by which TRs gain or lose multiple motifs at once, in contrast to indels; Fan and Chu, 2007) and unequal recombination (redistribution of motifs among gametes due to the misalignment of homologous TRs during cross-over; Smith, 1976). While replication slippage is key to the maintenance of TRs within populations (Ohta, 1983b; Stephan, 1987), unequal recombination tends to increase variation in copy number (Stephan, 1986). In turn, greater variation leads to more efficient selection against TR copies in the face of genetic drift (Stephan, 1987).

Here, we relax two assumptions made in all these models in order to gain greater insights into the processes that shape variation in TRs within and between populations. First, instead of assuming that individuals mate randomly, we will consider the common form of non-random mating that is partial selfing. Partial selfing leads to excess homozygosity, which in turn affects selection, drift and recombination (Burgarella and Glémin, 2017). This excess homozygosity should be especially relevant to TRs evolution as unequal recombination depends on the variance among homologous copies. Second, rather than assuming that selection on TRs length acts at the haploid gametic stage and is only purifying based on additive effects (i.e. each additional repeat has the same fitness effect, but see Stephan, 1987 for a model of truncation-selection), we consider various modes of selection acting at the diploid stage: (i) non-additive effects in TRs length, in line with the observation that in some cases the onset of diseases is determined on the excess of repeats beyond a threshold (Usdin et al., 2015; Depienne and Mandel, 2021); (ii) interactions between homologous TRs within individuals, motivated by studies showing that the lack of TR homology can compromise chromosomal stability (for instance by inducing chromosomal loops or supercoiling, Usdin et al., 2015; Verbiest et al., 2023); and (iii) stabilizing selection for an optimal TRs length, based on eQTL and mQTL studies that have demonstrated that some TRs can influence gene expression and regulation and therefore contribute to phenotypic variation (Quilez et al., 2016; Gymrek et al., 2016; Fotsing et al., 2019; Verbiest et al., 2023).

## 2 Model

### 2.1 Life-cycle, trait and its distribution

We consider a population of diploid hermaphrodites of constant size *N* with the following life-cycle (fig. 1A, table 1 for a list of symbols): (1) Each adult produces a large number of gametes according to its fecundity and then dies. (2) Gametes fuse together to form zygotes. With probability *α*, a zygote is produced by combining two gametes of the same individual, or with complementary probability 1 − *α*, of two different individuals, so that the parameter *α* is the selfing rate. (3) Zygotes compete randomly to form the *N* adults of the next generation.

**Table 1:**
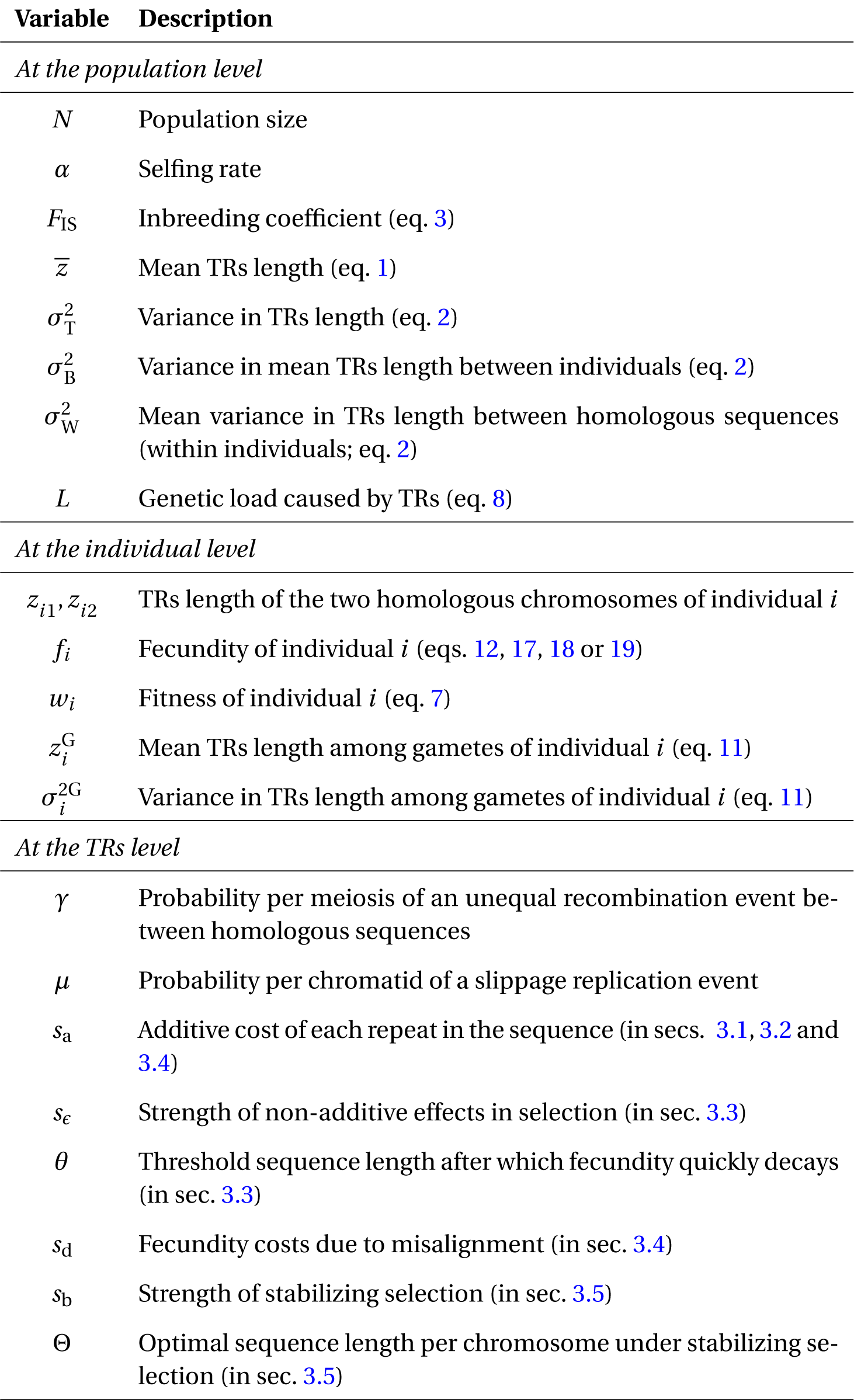
Summary of variables and parameters of the model.

**Figure 1:**
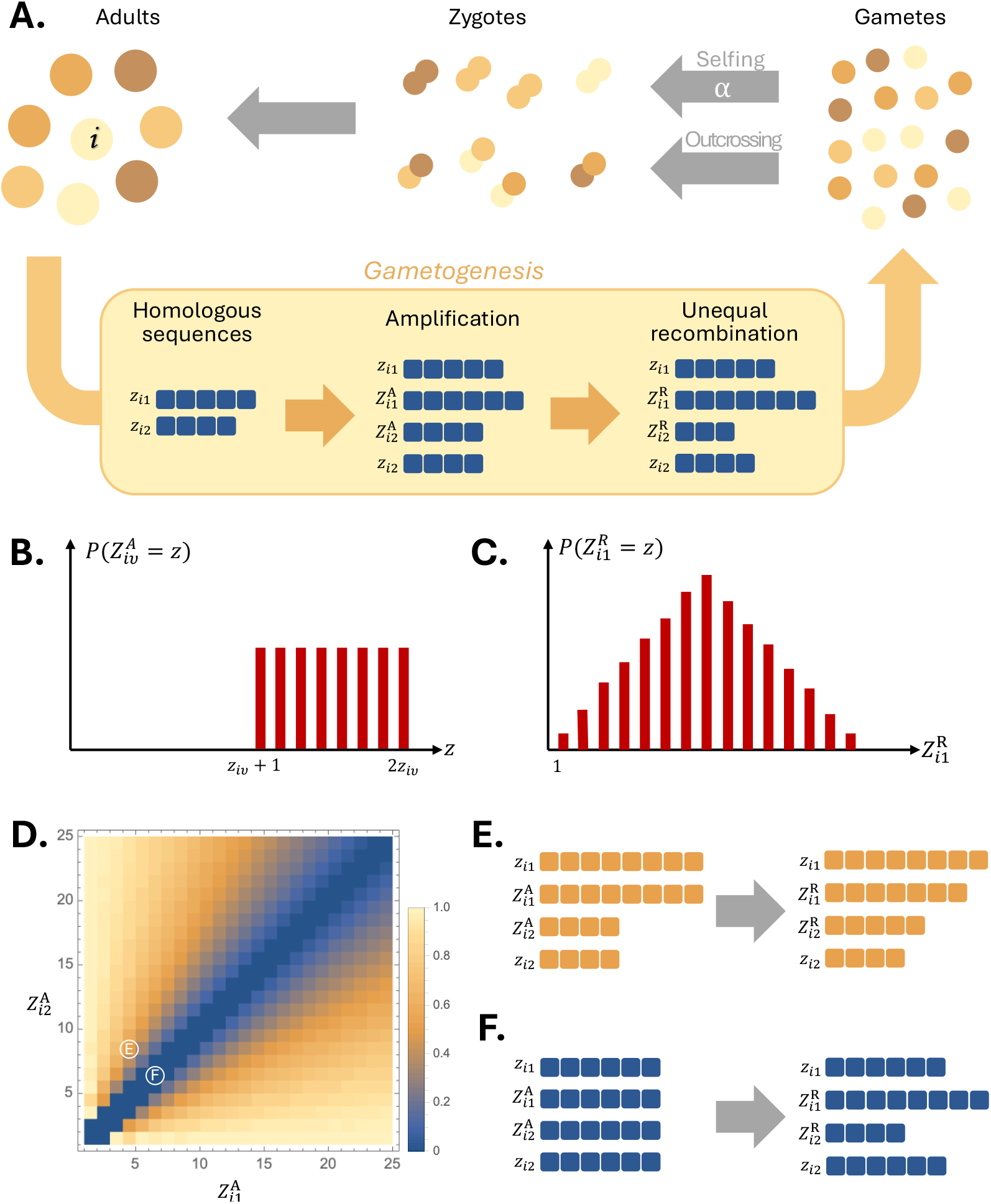
**A**. Life cycle described in sec. 2.1, with a highlighted example of the production of gametes of an adult *i*, as detailed in sec. 2.2. In this example *z*_*i*1_ *=* 5 and 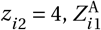 is amplified while *Z* ^A^ is not, and unequal recombination between 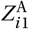 and 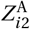 takes place. **B**. Probability distribution for the TRs length after amplification 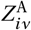 given template *z*_*iν*_ (eq. 9). **C**. Probability distribution for the TRs length after unequal recombination 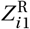 given amplified sequences 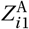 and 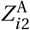 (eq. 10). **D**. Probability that unequal recombination decreases differences between gametes, i.e. probability that 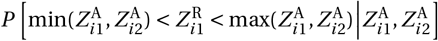. Examples of both cases, when unequal recombination (**E**.) decreases or (**F**.) increases variance between gametes.

Each individual *i* ∈ {1,…, *N*} is characterized by two positive integers, *z*_*i*1_≥ 1 and *z*_*i*2_≥ 1, which are the lengths of a TRs on the paternally and maternally-inherited chromosomes (fig. 1A for illustration). Our goal is to investigate the effect of selfing rate *α* on the evolution of this TRs. To do so, we will track the evolutionary dynamics of the population mean,

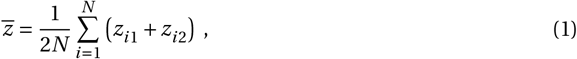

and variance,

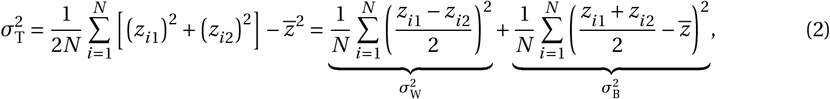

which we decomposed as the sum between the variance within 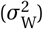 and between 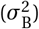 individuals. This decomposition of variance allows us to define the *F* -statistic (also referred to as the *R*-statistic in the context of microsatellites, see eq. (13) in Slatkin, 1995) as

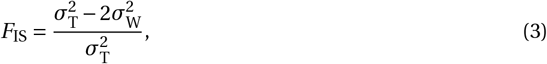

such that

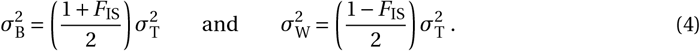

These equations reflect that under full outcrossing (*α =* 0), *F*_IS_ *=* 0 as the variance within individuals is equal to the variance between individuals (since there is no correlation between homologous sequences). Conversely, *F*_IS_ *=* 1 under full selfing (*α =* 1) and homozygosity is increased 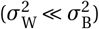.

In general, the expected change 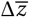 in mean TRs length 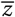 over one generation can be expressed as

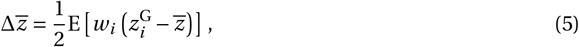

where 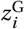 denotes the expected TRs length in a gamete of individual *i*, and *w*_*i*_ denotes the expected number of gametes of individual *i* that are recruited in the next generation. Similarly, the expected change in total variance 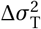 reads as

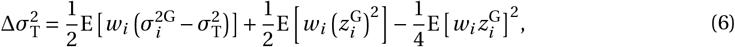

where 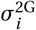 is the variance in sequence length among gametes of individual *i*. In both eqs. (5) and (6), the expectation E[·] is taken over all individuals *i* and all events that occur in a full iteration of the life-cycle (Appendices A.2 and A.3 for derivation of eqs. 5 and 6 respectively).

We assume that the fecundity of an individual depends on the TRs it carries, so that the fecundity *f*_*i*_ of individual *i* can be written as *f*_*i*_ *= f*(*z*_*i*1_, *z*_*i*2_). Given our assumptions on the life-cycle, the fitness *w*_*i*_ of this individual is given by

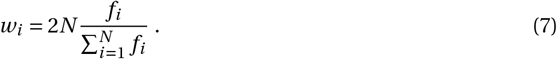

We’ll consider different forms for *f*_*i*_ to reflect different types of selection on TRs, and quantify the impact of TRs on the population by the genetic load

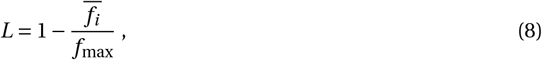

where 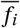 is the average fecundity in the population and *f*_max_ is the maximum fecundity calculated when extra repeats are absent.

In the next section, we characterize the moments 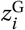 and 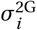 that appear in eqs. (5) and (6) as a result of meiosis and gametogenesis.

### 2.2 Meiosis & Gametogenesis

We assume that during meiosis and gametogenesis, three processes can influence the distribution of TRs among the recruited gametes of an individual: amplification, unequal recombination, and Mendelian segregation.

#### 2.2.1 Amplification due to replication slippage

First, the length of a TR sequence may be amplified during interphase due to replication slippage (e.g. fig. 1A; Levinson and Gutman, 1987; Charlesworth et al., 1994; Buschiazzo and Gemmell, 2006; see Khristich and Mirkin, 2020 for deeper discussion on mutational processes in TRs). To model this process, let 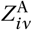 be a random variable for the length of the TRs in a focal germ cell after replication on the sister (inner) chromatid whose template is *z*_*iν*_ (with *ν* ∈ {1, 2}) in individual *i*. We assume that amplification takes place independently with probability *µ* during the replication of each new chromatid, in which case saltatory amplification occurs following the model of Stephan (1987). This model, which is based on the evidence that TRs length tends to grow due to mutation in a way that increases with the length of the parental sequence (Slatkin, 1995; Buschiazzo and Gemmell, 2006), assumes that the TRs length in the new inner chromatid 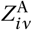 increases by a random number of TRs that is uniformly distributed between 1 and the template size *z*_*iν*_ (fig. 1B). That is, with probability *µ*, we have

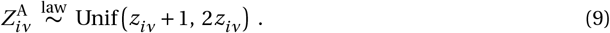

With complementary probability 1 −*µ*, no amplification happens so that the new sister chromatid is identical to its template, i.e. 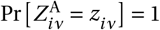.

#### 2.2.2. Unequal recombination

Following replication, the newly formed inner chromatids may recombine. When chromosome pairing during synapsis is correct, recombination does not affect the length of TRs. When however pairing is incorrect, which readily occurs owing to the repetitive nature of TRs, unequal crossover causes the number of TRs to be redistributed between chromatids during recombination (Krüger and Vogel, 1975; Perelson and Bell, 1977; Ohta, 1981; Ohta and Kimura, 1981; Stephan, 1986). To model this process, let us denote by 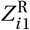 and 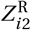 the random variables for the TR copy number produced by the inner chromatids after recombination. We assume that unequal recombination occurs with probability *γ*, in which case we have

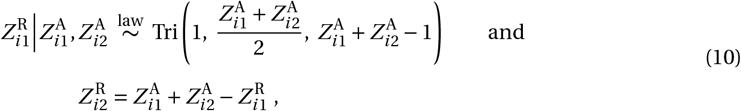

where Tri denotes the Triangular distribution (fig. 1C; Takahata, 1981; Stephan, 1986). This model assumes that any of the 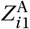 sites of the chromatid 1 is equally likely to cross over any of the 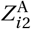 sites of its homologous chromatid during synapsis and that the total number of copies is conserved, i.e. 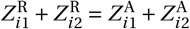 (Takahata, 1981; Stephan, 1986).

#### 2.2.3 Mendelian segregation

Finally, random Mendelian segregation distributes the products of meiosis fairly among the gametes of an individual. To see these effects, let us denote by 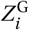 the random TRs length in the sequence of a gamete sampled among all gametes produced by individual *i*, whose TRs lengths are *z*_*i*1_and *z*_*i*2_. The moments of 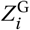 that are necessary to our analysis (i.e. that appear in eqs. 5 and 6) are then given by

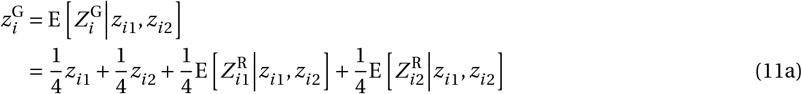

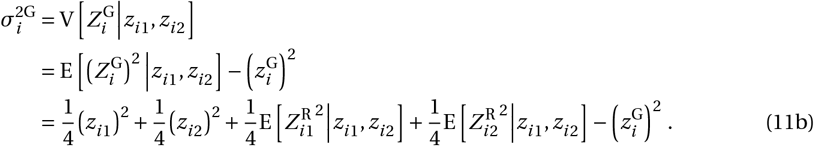

In both equations above, the first two terms correspond to the outer chromatids, while the next two terms represent the inner chromatids, which undergo amplification and unequal recombination.

We investigate the evolution of TRs under different selection regimes with two complementary approaches. We compute the moments in eq. (11) using the distributions given in eqs. (9) and (10) (details in Appendix A.4), and then substitute these moments into eqs. (5) and (6) to construct an analytical framework for the change in mean and variance within the population across generations. We also perform individual-based simulations using SLiM v4.3, considering the life cycle and gametogenesis described above, with further details of the implementation available in Appendix B (Haller and Messer, 2023). Our main results are summarized below.

## 3 Results

### 3.1 Shorter TR sequences under purifying selection in selfing populations

As a baseline, we assume that TRs are under purifying selection with additive effects of repeats, e.g. owing to the time and energy cells invest to replicate TRs in the genome (Stephan, 1986, 1987; Charlesworth et al., 1994; Buschiazzo and Gemmell, 2006; Verbiest et al., 2023). We assume that the fecundity *f*_*i*_ of a focal individual *i* carrying sequences of lengths *z*_*i* 1_ and *z*_*i* 2_ is

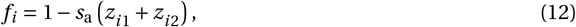

where *s*_a_ tunes the strength of purifying selection.

Because the expressions of the population mean and variance (eqs. 5 and 6) are too complicated to have an analytical form in general, we first assume that selection and amplification are weak (*s*_a_ ∼ 𝒪 (*δ*) and *µ* ∼ 𝒪 (*δ*) where *δ* is small parameter) and that population is large (*N* ∼ 𝒪 (1/*δ*)), obtaining

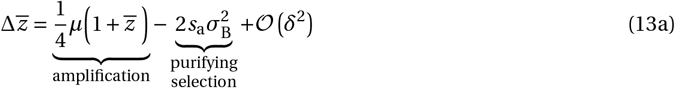

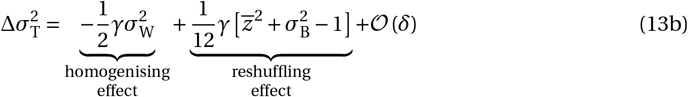

for the dynamics of the mean and variance in TRs length (Appendix A.6 for derivation). The two terms of eq. (13a) highlight how the change in TRs length depends on a balance between: (i) amplification, whose effects are proportional to the mean 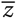 due to its saltatory nature (i.e. larger sequences gain on average more TR copies); and (ii) purifying selection, whose effects are proportional to the variance between individuals 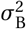 as selection takes place among adults. The two terms of eq. (13b), meanwhile, reflect how unequal recombination can either decrease or increase the variance in TRs length, depending on how the variance in TRs length is distributed within and between individuals (fig. 1D). If the variance within individuals 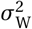 is large (compared to 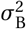), then the first term of eq. (13b), which is negative, tends to dominate, indicating that recombination tends to reduce total variance. This is because when recombination takes place among homologous TRs that have sufficiently different lengths, recombination tends to homogenise these (e.g. fig. 1E). If, however, the variance within individuals 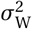 is small (compared to 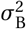), recombination in this case makes these sequences more different by reshuffling TRs, so that the variance increases (e.g. fig. 1F). This is captured by the second term of eq. (13b).

Comparing eqs. (13a) and (13b) shows that there is a separation of timescales between the dynamics of the mean and variance: changes in mean are of order *δ* while changes in the variance are of order 1 (under our assumption that *s*_a_ ∼ 𝒪 (*δ*) and *µ* ∼ 𝒪 (*δ*) while *γ* ∼ 𝒪 (1)). This entails that the dynamics of the variance should stabilise to an equilibrium 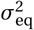 before the mean when *δ* is small. Solving 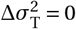 with eq. (4) for 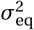 with a given 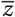, we obtain that this equilibrium is

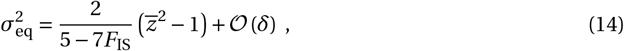

where

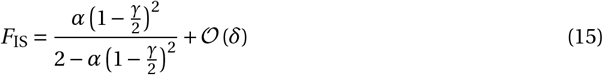

(Appendix A.5 for derivation). Equations (14) and (15) show that for a given population mean 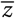, greater selfing rate *α* leads to a greater equilibrium variance. This is because selfing increases the reshuffling effect relative to the homogenizing effect via a decrease in 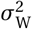 compared to 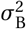 in eq. (13b). The effect of selfing on the inbreeding coefficient *F*_IS_ is reduced by unequal recombination because unequal recombination decreases homozygosity (this effect of *γ* is weighted by 1/2 in eq. 15 as only the inner chromatids can undergo unequal recombination).

Solving 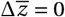 for 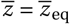 with 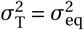 given by eq. (14) (and using eq. 4), we obtain that the mean of the equilibrium distribution of TRs length is

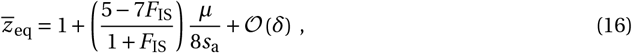

which shows that selfing reduces the mean TR copy number in the population (fig. 2A). This reduction is due to a greater proportion of the total variance in TRs that is between individuals, which increases the efficacy of selection (recall 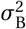 in eq. 13a). Plugging the mean in eq. (16) into eq. (14), we find that selfing also decreases the total variance among sequences in the population (fig. 2B). Because selfing leads to fewer TR copies, it also reduces the load associated with them (fig. 2C). Indeed, using eq. (8) with eqs. (12) and (16) we obtain the genetic load 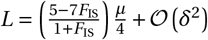. Therefore, genetic load decreases with selfing owing to shorter TRs (fig. 2C). How much selfing reduces the mean and variance in TRs length depends on the amplification-to-selection ratio *µ*/*s*_a_ (curves in fig. 2A).

**Figure 2:**
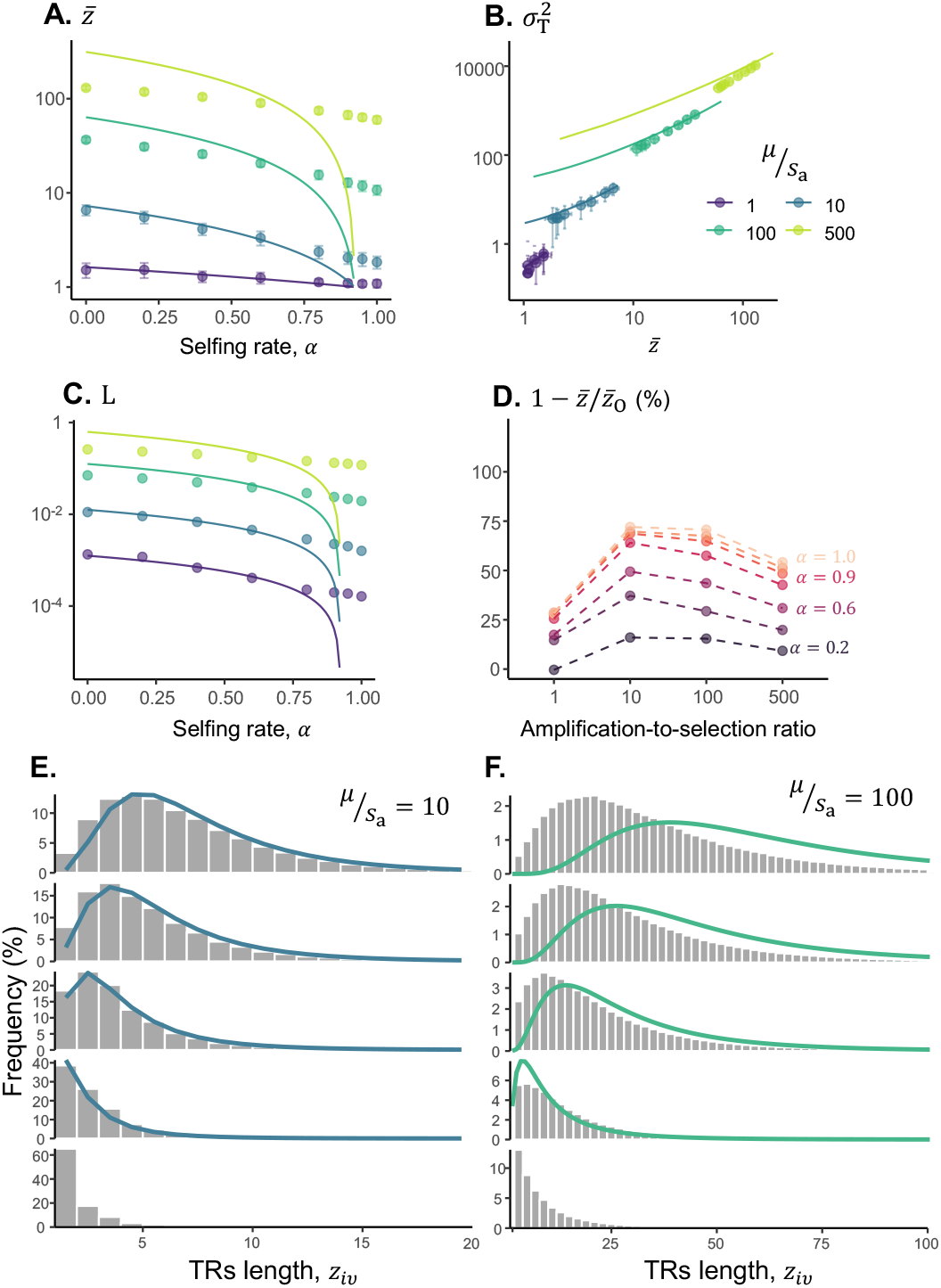
**A**. Mean TRs length 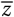 (in logscale) for various selfing rate *α*, with dots as averages from simulations (error bars show standard deviation; Appendix B for details on simulations). Lines represent the theoretical prediction obtained with eqs. (14) and (16). Parameters: *s*_a_ *=* 0.001, *γ =* 0.1, *N =* 2^*′*^000. **B**. Relationship between the total variance 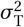 and the mean TRs length 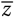 (both on logscale) for different selfing rates. Dots represent averages from simulations, while lines show theoretical predictions based on eqs.(14) and (16). Notably, eq.(14) accurately captures the relationship between the moments, even in cases where eq. (16) does not accurately predict the mean. Parameters: same as A. **C**. Genetic load *L* (in logscale) as a function of selfing rate *α*, with dots as averages from simulations using eq. (16). Lines represent the theoretical prediction. Parameters: same as A. **D**. Reduction in the mean TRs length 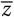 relative to outcrossing populations 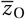 for different selfing rates (from bottom to top: *α =* 0.2, 0.4, 0.6, 0.8, 0.9, 0.95 and 1.0). **E**. and **F**. Distribution of TRs length in populations with different selfing rates (from top to bottom: *α =* 0, 0.25, 0.5, 0.75 and 1.0). Histograms are based on simulations and lines represent the theoretical prediction obtained with a lognormal distribution with mean given by eq. (16) and variance given by eq. (14). Parameters: *µ*/*s*_a_ *=* 10 in E and *µ*/*s*_a_ *=* 100 in F, other parameters as A.

We performed individual-based simulations to investigate the case *µ*/*s*_a_ ≫ 1 (points in fig. 2A). As predicted, our analytical approach for 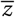 matches simulations quite well when amplification and selection are of similar order and both small relative to unequal recombination; with the exception when *F*_IS_ > 5/7 (*α* > 0.8). In this latter case, our analytical model predicts the complete loss of repeated motifs (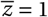 and 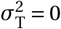) while simulations still show the maintenance of small TR sequences (mostly due to amplification events). This is because our separation of timescales breaks down when 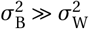 (see eq. 13b). When *µ*/*s*_a_ ≳ 100, our simulations show shorter TRs in outcrossing populations than predicted by eq. (16).

To see how the effects of selfing interact with other factors, it is useful to look at the relative reduction in copy number in a selfing population compared to an outcrossing one, all else being equal (i.e. we measured 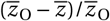 where 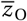 is the mean TRs length in outcrossing populations, *α =* 0). This shows that the relative reduction due to selfing is smaller in the regime where *µ* >> *s*_a_ (fig. 2D). This is because : (i) amplification increases TRs length similarly in outcrossing and selfing populations thereby reducing the differences they show (recall eq. 13a), and (ii) amplification decreases the excess homozygosity caused by selfing and thus mitigates its effect.

Simulations also reveal that the skewness and kurtosis of the TR copy number distribution in the population are greater when selfing is more frequent (fig. 2E and F for examples; Supplementary fig. S1). Altogether, this means that under partial selfing, we expect more outliers with long sequences in a particular sample compared to if TRs length were normally distributed. In fact, the distribution of TRs length shows a good fit to a lognormal distribution whose mean and variance are given by eqs. (14) and (16). This fit is particularly good when *s*_a_ *< µ < γ* (see solid lines in fig. 2E), but less good when *s*_a_ *< µ* ≈ *γ* (fig. 2F).

### 3.2 The effect of genetic drift

We used simulations to investigate the effect of genetic drift in small populations, in particular when 2*N* ≲ 1/*γ*. These show that for a given selfing rate, smaller populations tend to have longer TRs (compare points of same colour in fig. 3A). This is due to less efficient purifying selection in smaller populations, which leads to an accumulation of more repeats. In fact, the variances between and within individuals are both greater in smaller populations (fig. 3B for total variance 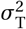, see Supplementary fig. S2 for each component). Additionally, the reduction in TRs length due to selfing is proportionally smaller in small populations (e.g., compare points when *N =* 10 in fig. 3A). This is presumably because small outcrossing populations show some level of inbreeding due to sampling effects and thus greater homozygosity.

**Figure 3:**
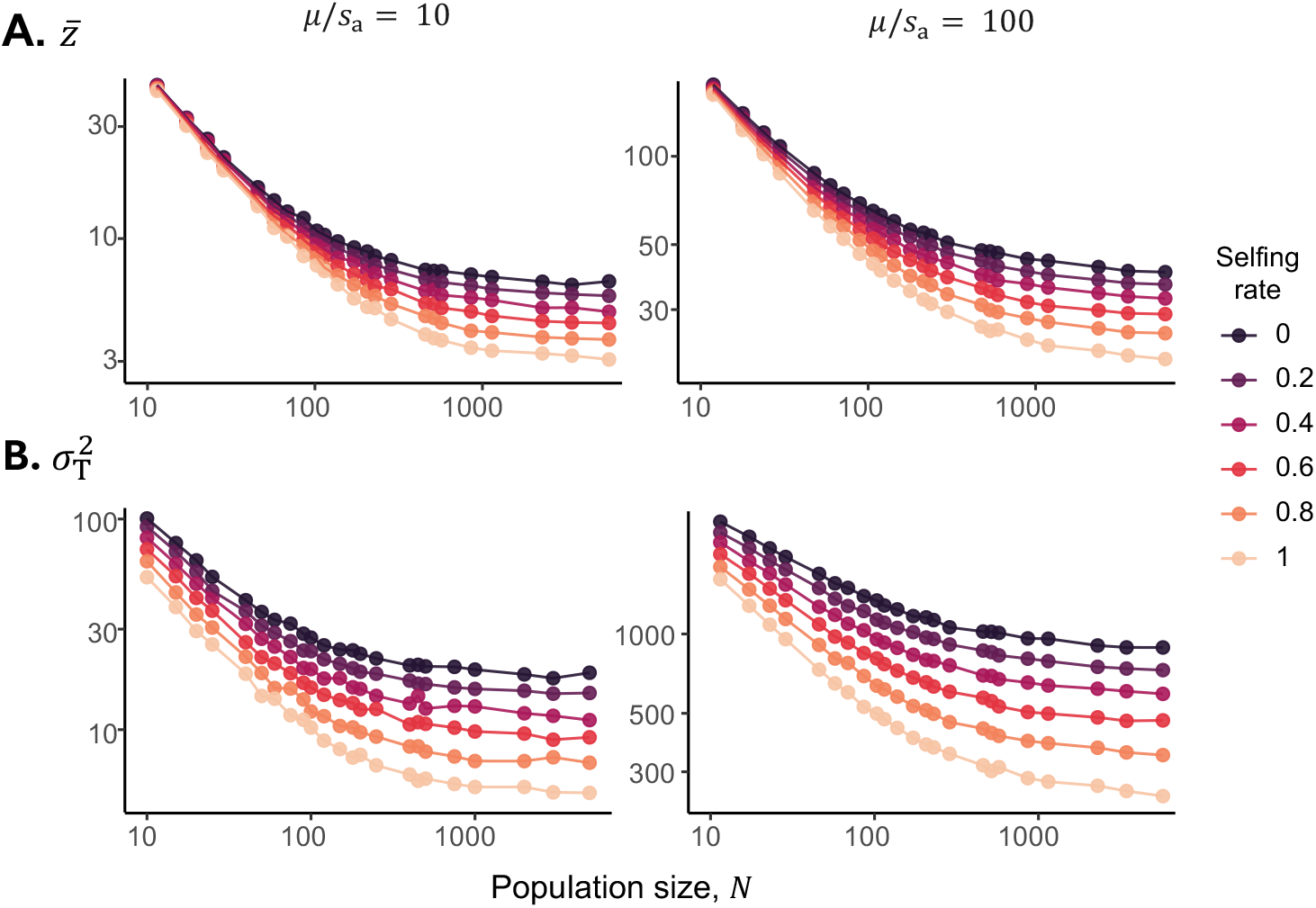
**A**. The mean TRs length 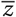 and **B**. total variance 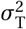 (both in logscales) in simulations with small and intermediate population sizes *N* (between 10 and 5000 individuals, in logscale) and for populations with various selfing rates (see legend). The number of replicates varies between population sizes, with at least 10 simulations for each parameter combination. Parameters: amplification-to-selection ratio *µ*/*s*_a_ *=* 10 (left) and *µ*/*s*_a_ *=* 100 (right column), other parameters: *γ =* 0.01, *s*_a_ *=* 0.001.

We also performed simulations with different population sizes *N* such that the effective population size *N*_e_ is held constant for different selfing rates *α* (with *N*_e_ *=* (1 −*α*/2)*N*, as in eq. 6 of Pollak, 1987; Supplementary fig. S3). As expected, these simulations show that the reduction due to selfing occurs when controlling for effective population size. More broadly, this shows that the effects of selfing via *N*_e_ are always weaker than those described in section 3.1 i.e., via selection (by boosting variance between individuals) and unequal recombination.

### 3.3 Truncation-like selection increases variance in TR copy number

We now consider the case where the TRs has non-additive effects such that the fitness of an individual rapidly declines after the number of repeats it carries goes beyond a certain threshold. Specifically, we assume that fecundity is

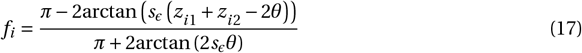

where 2*θ* is a threshold for the total number of TR, after which fecundity decreases towards zero at a rate that depends on the parameter *s*_*ϵ*_ > 0. When *s*_*ϵ*_ ≪ 1, eq. (17) approaches additive effects and we recover eq. (12) with a cost *s*_a_ → 2*s*_*ϵ*_/*π* per TR copy, regardless of *θ*. When *s*_*ϵ*_ ≫ 1, fecundity behaves similarly to a step function (in the limit *s*_*ϵ*_ → ∞): *f*_*i*_ *=* 1 for *z*_*i* 1_ *+ z*_*i* 2_ *<* 2*θ* and *f*_*i*_ *=* 0 for *z*_*i*1_*+ z*_*i*2_ > 2*θ* (this limiting case would be equivalent to truncation selection, which was studied in haploids under random mating in sec. 4 of Stephan, 1987).

Using individual-based simulations, we find that as the fecundity in eq. (17) approaches a truncation-selection curve (i.e., large *s*_*ϵ*_), the mean and variance in TRs both increase (fig. 4A). This is because selection against repeats when TRs are below the threshold (when *z*_*i* 1_ *+ z*_*i* 2_ *<* 2*θ*) is weaker when *s*_*ϵ*_ is large. In fact, as *s*_*ϵ*_ increases, the variance in TRs length per chromosome approaches (*θ* − 1)^2^ /12, which is the variance of a random variable following a uniform distribution between 1 and *θ* (this is also true when amplification increases, i.e. as *µ* gets large, in line with the results of Stephan, 1987, fig. 4B, Supplementary fig. S4 for more details on the TRs length distribution).

**Figure 4:**
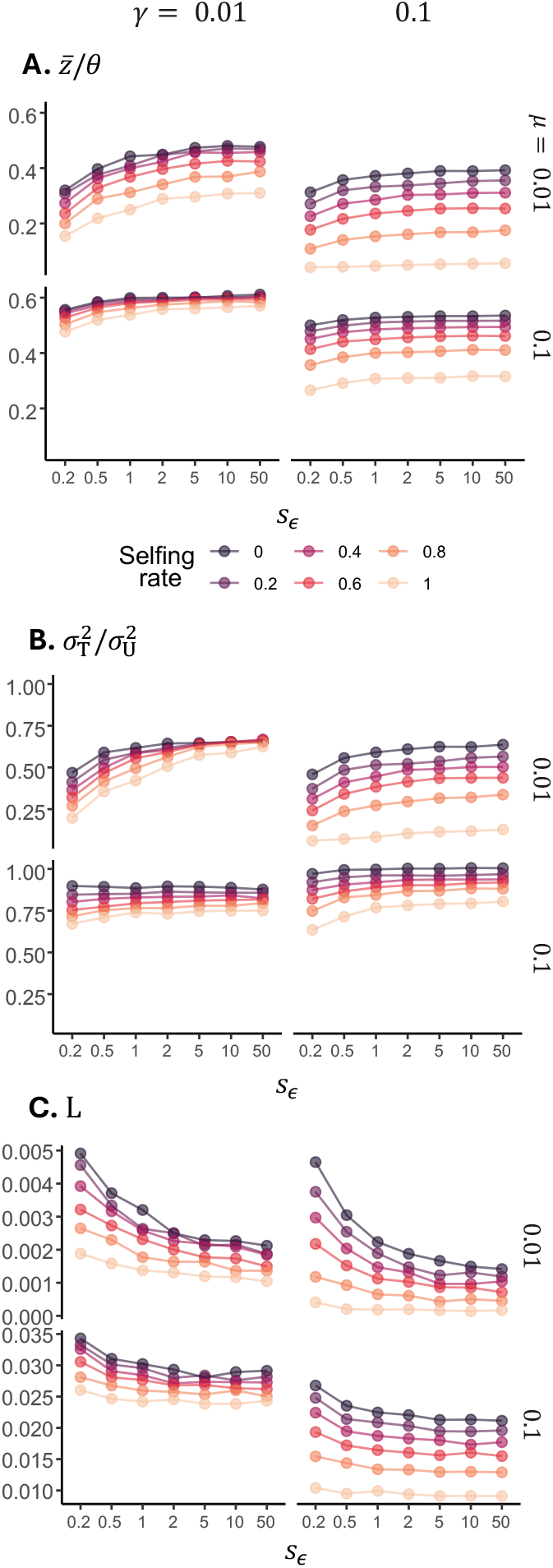
**A**. Mean TRs length 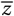 under non-additive fitness effects in eq. 17. Mean is represented as a fraction of the threshold length *θ*. Unequal recombination rate is *γ =* 0.01 in left column and *γ =* 0.1 in right column, while amplification rates are indicated in the right margin. Parameters: *N =* 2^*′*^000, *θ =* 100. **B**. Total variance in TRs length across various selfing rates (see legend in A) under non-additive fitness effects. As a reference, the variance is represented as a fraction of that in an uniform distribution where all sequences are equally present between 1 and the threshold *θ*, that is 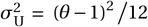. Parameters: same as A. **C**. Mean genetic load under non-additive fitness effects between repeats, calculated from the simulation with eq. (8) where *f*_max_ is given by eq. (17) with *z*_*i*1_ *= z*_*i*2_ *=* 1. Parameters: same as A.

Selfing meanwhile has similar effects to those found under additivity: it reduces the mean and variance in TRs length (compare dark and light lines in fig. 4A and B). This reduction, in turn, lowers the genetic load in the population (fig. 4C). The decrease in load is lower when selection is truncation-like (large *s*_*ϵ*_) since the deleteriousness of each repeat (as long as their total remains below 2*θ*) is lower than when effects are additive. The effects of amplification and unequal recombination on TRs length are also similar to those under additivity across all selfing rates: amplification tends to increase TRs while recombination tends to reduce them (as in haploid populations; Stephan, 1987).

### 3.4 Misalignment costs exacerbate the differences in TRs length of selfing and outcrossing populations

We now consider potential costs arising from the misalignment of homologous genes surrounding a TRs, e.g. when physical distortions during synapsis due to different TRs lengths create DNA secondary structures and loops that can lead to non-functional gametes after recombination (Balzano et al., 2021; Verbiest et al., 2023). To model this, let us assume that the fecundity of an individual is

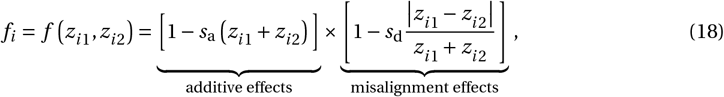

where the parameter *s*_d_ ≥ 0 tunes the cost of the misalignment. When *s*_d_ *=* 0, we recover the additive model (eq. 12), where each homologous TRs has effectively independent effects on fitness. When *s*_d_ is large, however, a difference in length between homologous TRs within individuals is also costly, reducing fitness. This can be seen as a form of under-dominance causing heterozygote disadvantage.

Our simulations show that in outcrossing populations, misalignment costs cause an increase in mean TRs length (dark points, especially *s*_d_ > 0.1, in fig. 5A). This is because any gamete carrying a TRs that is shorter than average will on average suffer a misalignment cost under random mating as it will likely fuse with a gamete carrying a longer TRs. Selection can therefore favour amplified sequences if it brings them closer in length to the average sequence, as long as *s*_a_ isn’t too large (fig. 5A with *s*_a_ *=* 0.001 fixed). Contrastingly, misalignment costs have weak to no effects in selfing populations since these show a deficit of heterozygosity (light points in fig. 5A). As a result, the difference in mean TRs length 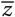 between outcrossing and selfing populations is greater when *s*_d_ is large (fig. 5A). The variance of TRs length are also affected by misalignment costs, but mostly in outcrossing populations where the variance within individuals 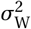 is reduced but between individuals 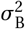 is increased. This is because misalignment costs simultaneously increase homozygosity and lead to longer sequences, inflating the total variance.

**Figure 5:**
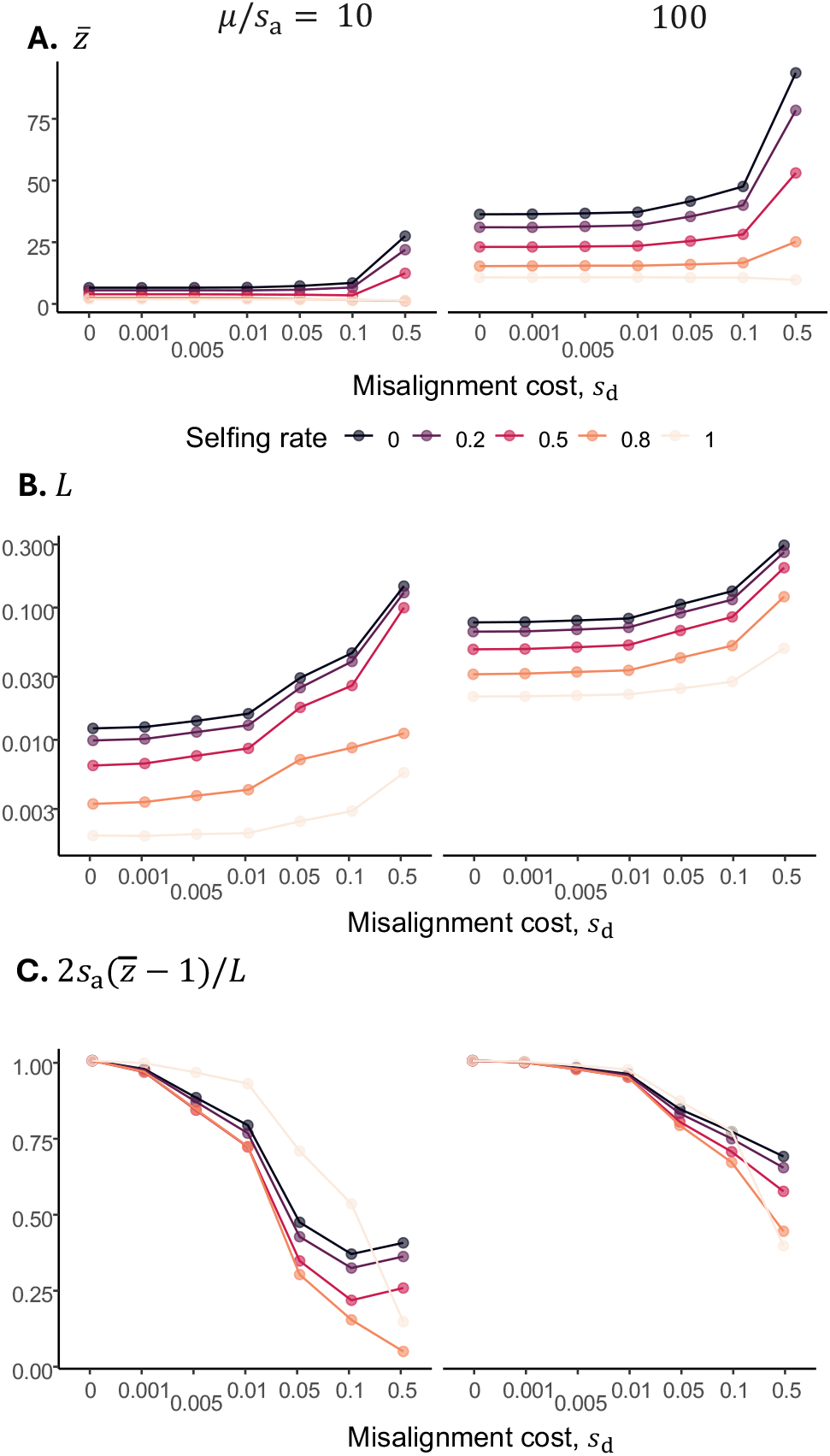
**A**. Mean TRs length 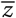 across various selfing rates (see legend) for different costs of misalignment (*s*_d_ in eq. 18 with *s*_a_ *=* 0.001 fixed). Parameters: *N =* 2^*′*^000, *γ =* 0.1. **B**. Genetic load *L* in the population (in logscale). Parameters: same as A. **C**. Proxy for the proportion of genetic load *L* due to additive effects. Parameters: same as A.

The effects of misalignment costs on the load *L* are similar to that on the mean TRs length, with the load increasing with misalignment costs in outcrossing but remaining similar in selfing populations (compare dark and light points in fig. 5B). To disentangle the contribution to the load of misalignment effects from an increase in 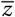, we computed 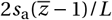 to measure the proportion of the load that is due to additive costs (i.e. arising from the first term in eq. 18). When *s*_d_ *=* 0, the entire load is due to additive effects, but when *s*_d_ is larger, the costs of misalignment represent a larger fraction of the load despite longer sequences (fig. 5C). In other words, misalignment costs grow faster than additive costs as *s*_d_ increases, and this is especially true in outcrossing populations (due to deficit of heterozygotes in selfing populations).

### 3.5 Partial selfing reduces load when TRs length is under stabilising selection

Finally, we examine the case where TRs are under stabilizing selection for an optimal length, which aligns with recent evidence suggesting that some TRs can play functional roles in different biological processes (Balzano et al., 2021 for examples) and contribute to phenotypic variation (Quilez et al., 2016; Gymrek et al., 2016; Fotsing et al., 2019; Verbiest et al., 2023). We therefore assume fecundity is given by

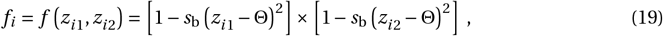

where the parameter *s*_b_ > 0 tunes the strength of stabilizing selection for the optimum Θ. We show in Appendix A.6.3 that by plugging eq. (19) into eq. (5), we obtain that the mean at mutation-selection balance in a large population can be expressed as

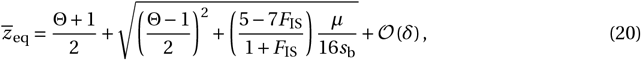

where we assumed that distribution of TRs length is approximately normal (which here is justified by stabilising selection; Walsh and Lynch, 2018). With weak selection, the population variance in TRs length is given still by eq. (14). The load meanwhile can be approximated as 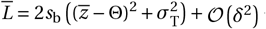 using eq. (19) in eq. (8) with *f*_max_ *=* 1. We also performed individual-based simulations. These, together with eq. (20), reveal that the effects of selfing depend on how large the optimal length Θ is compared to the amplification-to-selection ratio *µ/s*_b_ (fig. 6).

**Figure 6:**
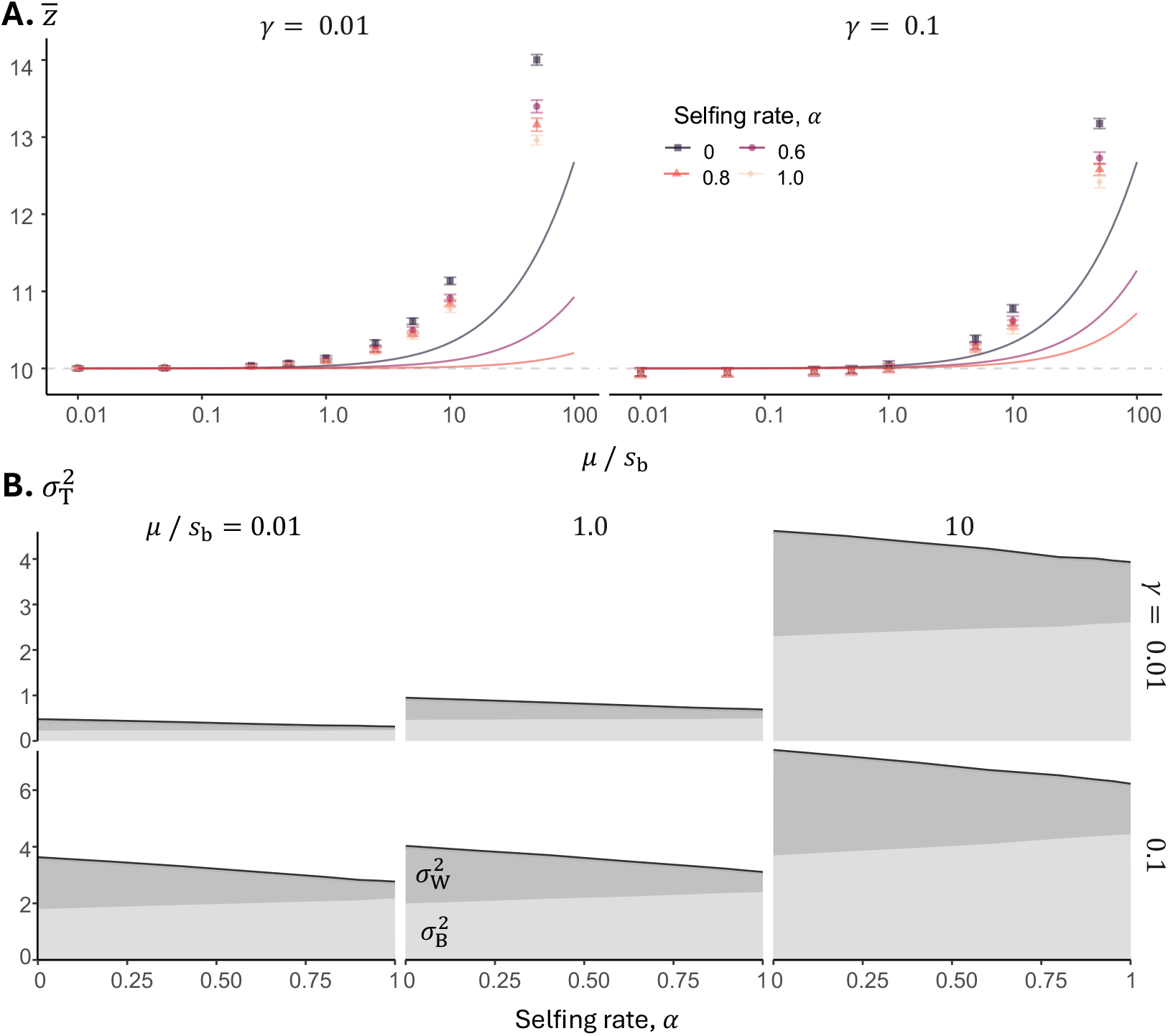
**A**. Mean TRs length 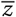 across different values of amplification-to-selection ratio (*µ*/*s*_b_) for various selfing rates (see legend). Unequal recombination rate is *γ =* 0.01 (left) and *γ =* 0.1 (right column). Curves present the analytical results of eq. (20). The horizontal gray line indicates Θ *=* 10. Parameters: *s*_b_ *=* 0.005, *N =* 5^*′*^000. **B**. Variance in TRs length across various selfing rates under stabilizing selection (from left to right, *µ*/*s*_b_ *=* 0.01, 1 and 10). Top row has *γ =* 0.01, while *γ =* 0.1 in bottom row. Shades of gray under the curve represent the proportion of variation between (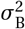, light gray) and within individuals (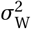, dark gray). Parameters: same as A.

When Θ ≫ *µ*/*s*_b_, the mean TRs length 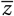 is close to Θ, regardless of selfing rate. This can be seen from eq. (20) putting *µ*/*s*_b_ to zero (see also fig. 6A for simulations). This is because stabilizing selection here is strong and amplification events are rare so that rare mutations away from the optimum are efficiently purged in both outcrossing and selfing populations. In fact, the variance around Θ in the population is always small, especially so when unequal recombination is infrequent (compare top and bottom plots in left column, *µ*/*s*_b_ *=* 0.01, in fig. 6B). As a consequence of the absence of additional repeats and little variation, the load in the population is negligible (purple line in Supplementary fig. S5).

When Θ ≲ *µ*/*s*_b_, the mean TRs length in the population is greater than the optimum Θ as here stabilizing selection is weak and amplification events frequent (fig. 6A). The increase in 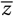 is smaller under selfing (compare points of different colours in fig. 6A) as selfing increases the proportion of variance between individuals increasing the efficacy of stabilizing selection (light gray area in fig. 6B).

Note that eq. 20 tends to underestimate 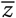 observed in simulations (fig. 6A). This is because eq. (20) neglects the loss of variation caused by stabilizing selection itself and thereby overestimates the efficiency of selection towards Θ. The reduction in both the mean and total variance in TRs length under selfing leads to a corresponding reduction in genetic load (Supplementary fig. S5).

## 4 Discussion

We identified two main pathways through which selfing influences TRs owing to excess homozygosity: (i) by increasing the effect that unequal recombination has on generating variation among sequences within individuals, and (ii) by increasing the variance in TRs between individuals. As a result of these effects, selection (here assumed to take place at the diploid stage) tends to be more efficient under partial selfing, leading to on average shorter and less diverse TRs. In turn, this means that selfing populations show lower genetic load due to TRs. These effects of selfing are especially strong in large populations.

### 4.1 Empirical implications

One implication of our results is that, all else being equal, TRs length should negatively correlate with selfing rates. More broadly, since the effects of selfing operate through increased homozygosity, any mechanism increasing homozygosity is expected to similarly reduce TRs length. Thus, TRs length should generally show a negative correlation with homozygosity, irrespective of the specific cause.

Genomic data to test these predictions remain sparse, primarily because TRs lengths within populations are rarely reported. In fact, TRs are frequently filtered out during early stages of genomic data processing. One relevant study is Viard et al. (1996), who investigated polymorphisms at four TRs across populations of partially selfing freshwater snails. They found substantial variation within populations (e.g., some individuals had twice as many repeats as others for a given sequence). However, no correlation between TRs length and inbreeding coefficients was observed, perhaps due to the limited variation in selfing rates across populations (all populations had *F*_IS_ > 0.7). Similarly, Zenke et al. (2011) found low variation in TRs length across highly inbred dog breeds, resulting in no detectable correlation between genome-wide inbreeding levels and TRs length.

Indirect evidence could potentially be drawn from cross-species comparisons. For example, humans have significantly longer homologous TRs than other primates (Rubinsztein et al., 1995), possibly due to their lower levels of homozygosity (Kuderna et al., 2023, but see Prado-Martinez et al., 2013 for differences in demographic histories that can also impact homozygosity). Additionally, genome-size comparisons across species frequently report smaller genomes in selfing species (Wright et al., 2008). Given that TRs typically constitute a conserved fraction (∼ 3%) of genomes (Srivastava et al., 2019), this trend might indirectly support our prediction. However, multiple factors influence genome size, making such indirect comparisons challenging. Further genomic analyses explicitly comparing TR variation across species or populations differing in mating systems or inbreeding levels are needed to rigorously test the predictions of our model.

### 4.2 The nature of selection on TRs

TRs are increasingly recognised for their potential role in complex traits, as highlighted by recent genome-wide association studies (reviewed in Depienne and Mandel, 2021 for humans and Verbiest et al., 2023 across various taxa), implying they likely have relevant but potentially non-straightforward fitness consequences. However, previous theoretical work on TRs evolution has primarily considered purifying selection in haploid populations (Stephan, 1987). Our model extends these analyses by explicitly incorporating diploidy, partial selfing, and by introducing two additional selective scenarios: (i) selection against misalignment costs (heterozygote disadvantage), and (ii) stabilising selection favouring an intermediate TRs length.

Our analyses revealed that selfing consistently reduces the genetic load associated with TRs across all selection regimes. The robustness of this finding implies that distinguishing among different selection types solely based on correlations between mean TRs length and inbreeding coefficients (*F*_IS_) may not be particularly informative. Nevertheless, the absence of such a correlation might indicate either strong stabilising selection or a mutation-driven equilibrium unaffected by mating system. This mutation-driven equilibrium is a balance between mutational processes like slippage, point mutation, expansion and contraction that lead to a stable mean TRs length (Kruglyak et al., 1998, 2000). Conversely, observing a very strong correlation could suggest selection against misalignment costs, as this scenario produced the largest differences in TRs length between selfing and outcrossing populations.

Under purifying selection, the distribution of TRs lengths is heavily skewed and characterised by frequent outliers, particularly when mean TRs lengths are short (under low amplification or high selfing rates). Our results suggest these distributions are well-described by a lognormal distribution, predicting many short sequences alongside occasional but readily sampled long sequences. This finding aligns with empirical observations of TRs frequently described as hypervariable (Jeffreys et al., 1985; Lareu et al., 1998; Legendre et al., 2007; Wei et al., 2014; Verbiest et al., 2023; Lundström et al., 2023). Additionally, our model predicts greater variation in TRs length within populations as mean TRs length increases, consistent with human data (Legendre et al., 2007; Duitama et al., 2014).

In the scenario of strong truncation-like selection, the average TRs length in the population remains substantially below the threshold above which it becomes deleterious (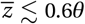 in fig. 4A). Selection thus minimises the risk of deleterious expansions through replication slippage or recombination within a lineage. Interestingly, empirical observations from medical genetics are consistent with this. For instance, Fragile X syndrome manifests beyond 200 repeats of a CGG motif, yet most human individuals carry only 5 to 40 repeats (Depienne and Mandel, 2021; Lundström et al., 2023).

### 4.3 Contrasts and limitations

Because selfing reduces effective population size and decreases effective recombination, it is typically associated with greater genetic load, except when load is driven by recessive deleterious mutations (Glémin, 2007; Hartfield and Glémin, 2014; Abu Awad and Roze, 2018; Sianta et al., 2023; Stetsenko and Roze, 2022; see Crow and Kimura, 1970 p. 299 for overview on mutation load, and Burgarella and Glémin, 2017 for review on effects of selfing). In such cases, selfing can reduce load by exposing these mutations to selection. In contrast, our analyses shows that, due to the unique way mutation and recombination affects TRs variation (saltatory amplification and unequal recombination), selfing leads to shorter TRs and lower genetic load, even when TR copies have additive fitness effects within each sequence and between homologous sequences (eq. 12).

Finally, our model is based on several simplifying assumptions. First, we considered a single well-mixed population. However, since TRs often contribute to genetic differentiation between populations (Slatkin, 1995; Goldstein et al., 1995), investigating TRs evolution in subdivided populations connected by limited dispersal could be relevant, especially as kin selection and inbreeding may interact there (Rousset, 2004). Second, we considered selection acting on a single TRs whereas in reality, each TRs exists within a genetic context that can also be affected by selfing (e.g. linkage with recessive deleterious mutations). Third, we assumed that replication slippage always increases TRs length at a constant rate. However, replication slippage can either increase or decrease TRs length depending on which DNA strand loops out, through there is a bias towards increases due to the flexibility of the newly synthesised strand (Knox et al., 2024). Additionally, we neglected other types of mutations, such as point mutations, as well as possible interactions between mutations (Kruglyak et al., 1998; Ellegren, 2002). The omission of point mutations is partially justified by evidence that replication slippage events occur 10–100 times more frequently than point mutations in TRs (e.g., in primates; Pumpernik et al., 2008). Fourth, we assumed that homologous TRs of a selfed individual are independent. While this assumption is appropriate where offspring arise from two gametes produced by separate meioses, other forms of reproduction, such as asexual reproduction by parthenogenesis, involve offspring deriving from gametes produced by the same meiosis. In these cases, homologous TRs lengths within offspring could be correlated. Fifth, we considered the evolution of TRs on autosomes although TRs are also abundant in sex chromosomes (e.g. TRs constitute about half of the Y chromosome in humans; Cooke, 1976). Under our assumptions, the non-recombining heterogametic chromosome (Y or W) would presumably accumulate repeats, due to its inability to purge repeats combined with biased replication slippage towards increased lengths. In contrast, evolution on the recombining homogametic chromosome (X or Z) would mirror that of an autosome, but with appropriately rescaled effective population size and unequal recombination rates.

### 4.4 Conclusions

Our results show that mating systems, specifically partial selfing and the associated increase in homozygosity, influence the evolutionary dynamics of TRs. Selfing consistently reduces TRs length and associated genetic load across multiple selection scenarios. This reduction is primarily due to the way unequal recombination generates TR variation within individuals and how this interacts with selfing. Empirical analyses comparing TR variation across populations or species differing in mating systems or inbreeding levels are needed to test these predictions rigorously.

## Data Availability Statement

The code used for our simulations is available here: https://vsudbrack.github.io/TandemRepeatsSM. Otherwise, the authors affirm that all data necessary for confirming the conclusions of the article are present within the article, figures, and tables.

## Acknowledgements

We thank Tanja Schwander for motivating this study, Ehouarn Le Faou for useful feedback, and William Toubiana for interesting discussions.

## Appendices

### A. Mathematical model

In this Appendix, we develop the mathematical framework describing the evolution of TRs length in a diploid, partially selfing population. In particular, we derive eqs. (5), (6), (14), (16), and (20) of the main text.

#### A.1 Moments of TRs length distribution

Let us first introduce some short hand notation. We denote the *n*-th moment of the distribution of TRs length as

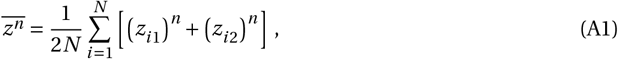

such that the mean TRs length is 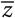 and the total variance 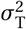 is

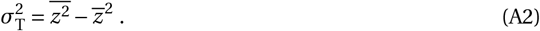

Additionally, we denote the second-order mixed moment between the two homologous chromosomes as

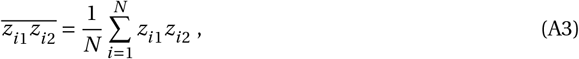

which allows us to write the covariance as

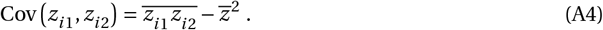

From these definitions above, together with eq. (2), it follows that

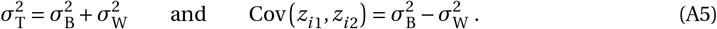

Finally, we note that because the labelling of the chromosomes (1 or 2) is arbitrary, we consider henceforth the distributions of TRs length in both homologous chromosomes to be equivalent, i.e. they share all expectations E[(*z*_*i*1_)^*n*^]= E[(*z*_*i*2_)^*n*^] for any *n* positive integer.

Unless other conditions are explicitly stated, we simplify our notations from now on by omitting in expectations, variances and covariances the conditions on values of 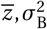 and 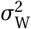, i.e. 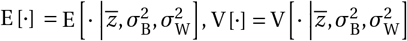 and 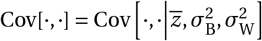.

#### A.2 Change in mean TRs length

Here, we derive eq. (5) of the main text by calculating the expected change in the mean number of copies 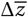 from one generation to the next in an infinitely large population. By definition, we have

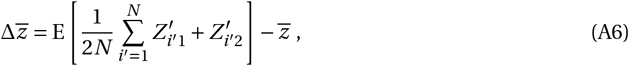

where *i*^*′*^ is used to index an individual in the next generation, whose maternal and paternal sequences contain 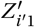 and 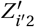 repeats respectively. We relate these sequences in the next generation with the gametes of the current generation, and thus 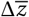 is given by

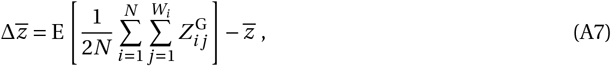

with 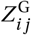 denoting the number of repeats in the TR sequence of the *j* -th recruited gamete of individual *i* in the current generation, and *W*_*i*_ denotes the total number of gametes recruited by individual *i* in the next generation. In order to simplify the sum, we condition our expectation on *W*_*i*_ to write

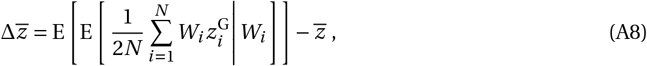

where 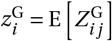 is the expected number of repeats in the TR sequence of a gamete of individual *i* (all *W*_*i*_ gametes of an individual are produced independently). Finally, we can calculate the expectation over *W*_*i*_, which is independent of 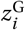,

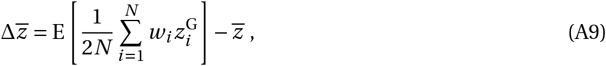

where *w*_*i*_ *=* E[*W*_*i*_], the expected number of gametes recruited by individual *i*. By switching the order of the expectation and the sum, we end up with

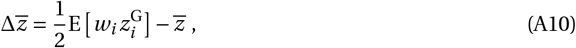

which becomes eq. (5) in the main text using the fact that E[*w*_*i*_] *=* 2 for a diploid population of constant size.

#### A.3 Change in variance in TRs length

We also track the change in total variance 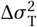 in an infinitely large population, which is defined as

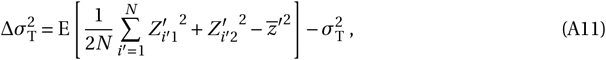

with the same notation as in eq. (A6), and 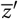 denoting the mean number of TR copies in the population in the next generation. Once again, we can write sequences in the next generation as gametes in the current generation, yielding

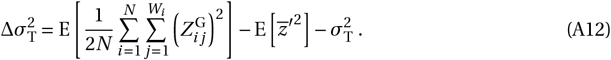

The second term of this expression is related to our previous section since we can write 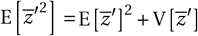, and in an infinite population 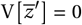, and thus 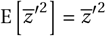. For the first term of eq. (A12), we can further simplify the sum by conditioning on *W*_*i*_, similarly to the previous section. This results in

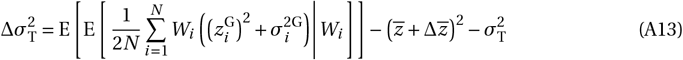

where 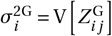 is the variance among gametes of individual *i*. Using eq. (A10) and switching the order of sum and expectation, we end up with

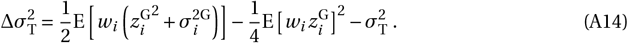

After rearranging the terms and using the fact that E[*w*_*i*_] *=* 2 in a population of constant size, we arrive at eq. (6) of the main text.

### A.4 Expectation and variance among gametes

#### A.4.1 Expectation and variance among gametes after amplification, 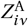

The expectation and variance of TRs length 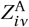 in the sister chromatid produced from a template with *z*_*iν*_ copies distributed according the uniform distribution of eq. (9) can be calculated directly from the sum of the different moments of the distribution. The expected value of the amplified sequence is given by

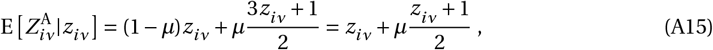

and the variance, computed with 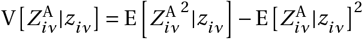, is given by

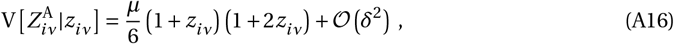

where *δ* is the order of the amplification rate *µ* ∼ 𝒪(*µ*).

#### A.4.2. Expectation and variance among gametes after recombination, 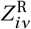

As a result of the distribution of eq. (10), given 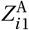 and 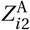, the expected number of TR copies in each inner chromatid after unequal crossover is given by

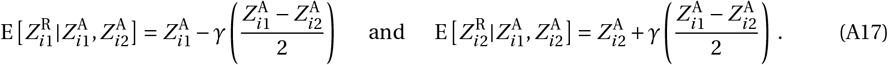

The expressions for expectation of 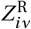 in eq. (A17) depends on the amplified lengths 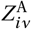, which are random variables themselves with expectation given by eq. (A15). To consider both steps altogether, we marginalize across all possible values of 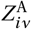. The expected number 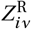 of TR copies in the inner chromatid after amplification and recombination given the number of copies in the parental adult is thus

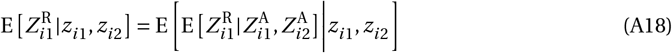

which can be calculated using the distribution in eq. (10) and eqs. (A15) and (A16), resulting in

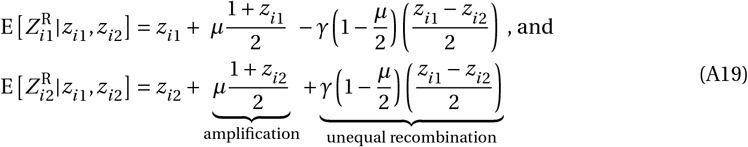

where we have labeled the terms capturing the changes due to the amplification of the focal inner chromatid and due to the unequal recombination between inner-chromatids, including the interaction between these two processes (e.g., an increase due to amplification of the homologous sequence followed by unequal recombination). The opposites signs of the “unequal recombination” term in eq. (A19) shows that unequal recombination tends to increase the number of TRs in the shorter sequence while decreasing the number of TRs in the longer sequence.

The variance in TRs length after recombination among gametes, eq. (10), depends on the parity of the total number of repeats after amplification. If the total number of repeats 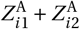 is even, then the center 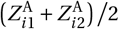 is also the mode of the distribution. If 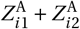 is odd, however, then the center 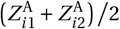 is actually the mean between the two most likely values of the distribution, 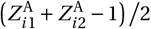 and 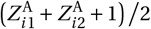. As a consequence, the variance in TRs length after recombination is slightly smaller when the total number of repeats is even,

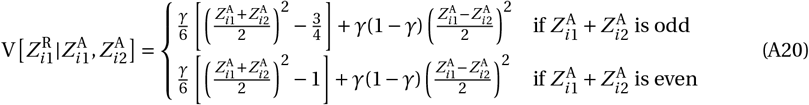

but the differences becomes increasingly negligible as sequences increase in length. For simplicity, we will assume that the sequences in the stationary regime are long enough such that the difference between the variances of an odd or even number of copies can be neglected and the variance is given by the latter. The expression for 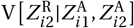 is also given by eq. (A20).

Similar to the previous calculation, the expressions for variance of 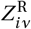 in eq. (A20) depends on the amplified sequence sizes 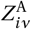 which are random variables themselves. We thus proceed to marginalize across all possible values of 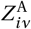. By doing so, the variance in number of copies after amplification and recombination given the number of copies in the parental adult is

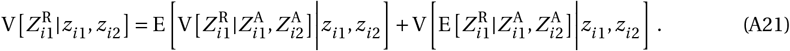

The first term is calculated using eq. (A20):

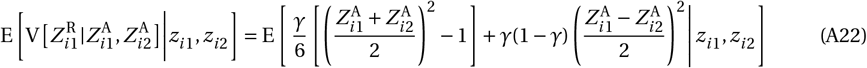

using 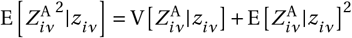 and the fact that amplification happens independently in each homologous chromosome, we can plug eqs. (A15) and (A16):

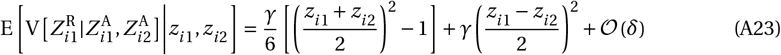

The second term of eq. (A21) is given by

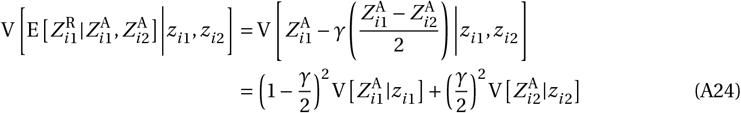

where we can use eq. (A16), yielding

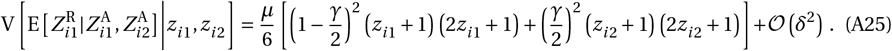

Putting eqs. (A23) and (A25) together into eq. (A21), we find that the variance between recombinant gametes is

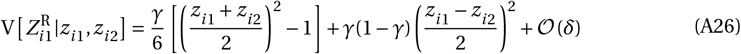

and 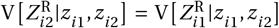. Amplification also tends to increase the variance among the inner chromatids, but it is neglected in eq. (A26) compared to unequal recombination of order 1, that is *γ* ∼ 𝒪(1) while *µ* ∼ 𝒪(*δ*).

### A.5 *F* -statistics

From the definition of *F*_IS_ in eq. (4) and eqs. (A5), we also find that

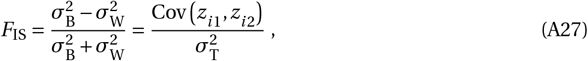

which means that in order to understand the changes in *F*_IS_, we have to calculate how the correlation between homologous chromosomes changes. Because of eq. (A3), we are thus interested in calculating

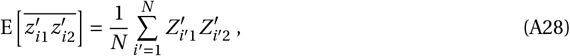

(same notation as eq. A6) and because the sequences in the next generation are the gametes of the current generation, we have

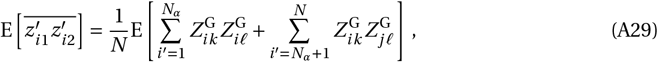

where *N*_*α*_ is the number of selfed individuals in the next generation, and index *k* and *𝓁* represent a random gamete sampled from individuals *i* or *j*. We can condition on knowing the number of selfed offspring, and switch the order of the expectation and the sums, yielding

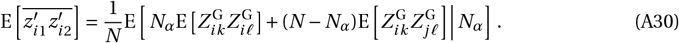

For the first term in this equation, we explicitly combine different gametes of individual *i* and take into account that unequal recombination is the main reason homologies are broken, that is

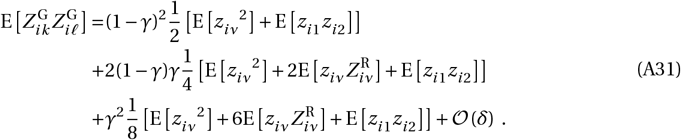

To simplify the terms with 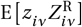, we notice that after an recombination event, the recombined gamete and the original gametes are independent. The calculation above simplifies to 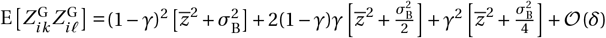.

The second term in eq. (A30) simplifies to

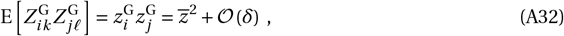

since gametes of different individuals are independent.

Putting these two terms into eq. (A30),

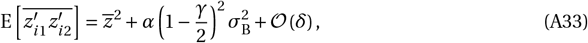

and thus back to the covariance using eq. (A3), we have

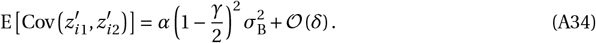

By using eqs. (A34) and (4) into eq. (A27), we have eq. (15) of the main text.

In the absence of unequal recombination (*γ =* 0), eq. (15) reduces to *F*_IS_ *= α/*(2 −*α*), which is the classical expression for the inbreeding coefficient under partial selfing (Wright, 1949; Pollak, 1987; Caballero and Hill, 1992). When unequal recombination is present, *γ* > 0, eq. (15) captures the fact that recombination breaks the associations among sister chromatids, generating heterozygosity even in selfers (that is to say, a zygote formed of *z*_*iν*_ and 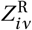 is no longer homozygous when recombination takes place). Because the outer chromatids do not undergo unequal recombination, the effect of recombination is weighted by 1/2 in eq. (15).

### A.6 Expected changes

#### A.6.1 Expected change in second moments of TRs length distribution

We start by calculating the expected change in the variance in TRs length. Under neutrality, i.e. when *w*_*i*_ *=* 2 *+*𝒪(*δ*) for each individual *i* in eq. (6), we obtain

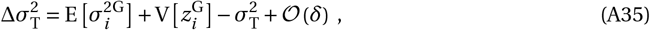

where recall that the expectation and variance are across individuals in the current generation given a 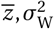 and 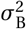. The first term of eq. (A35) is obtained by using eq. (11), yielding

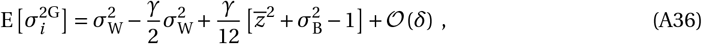

while the second term is

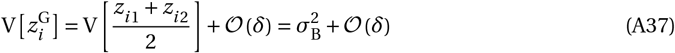

using eq. (11). Plugging these two results into eq. (A35), we have eq. (13b) of the main text.

#### A.6.2 Expected change in mean TRs length under purifying selection

Using eq. (7) and eq. (12), we find that the fitness *w*_*i*_ of the focal individual *i* is

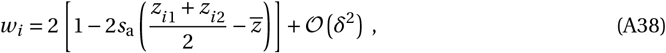

where *δ* is a parameter of order 𝒪(*s*_a_).

The expected change in the mean number of copies is found by plugging eqs. (A38) and (11) into eq. (5)

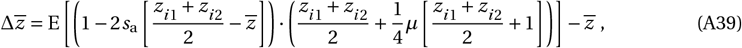

which simplifies up to first order of *δ* (recall *µ* ∼ 𝒪 (*δ*) and *s*_a_ ∼ 𝒪(*δ*)) to

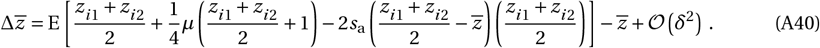

Finally, using the moments of the TRs length distribution to have the expectations, 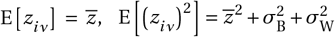 and 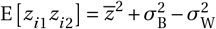, we end up with eq. (13a) of the main text.

#### A.6.3 Expected change in mean TRs length under stabilizing selection

We start by plugging *f*_*i*_ from eq. (19) into eq. (7), which yields after taking the average within the population

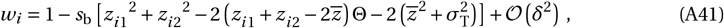

to the first order of selection (*s*_b_ ∼ 𝒪(*δ*)). By plugging eq. (A41) into eq. (5), we find that the change in mean, 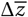, is given by

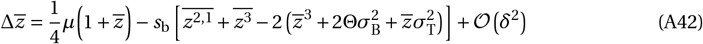

where the third moment of the TRs length distribution is denoted by 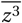 (eq. A1 with *n =* 3) and by 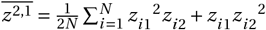. We will denote this third moment of the TRs length distribution as *κ*,

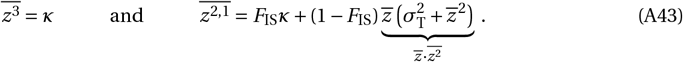

since *F*_IS_ is a probability of identity-by-descend, and this with probability *F*_IS_ the two sequences are equal length *z*_*i*1_ *= z*_*i*2_, and otherwise the sequences are independent. Plugging these third moments into (A42) produces

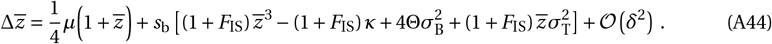

To proceed, we need an expression for the third moment *κ*. If we suppose that under stabilizing selection the TRs length is close to being normally distributed, then the third moment is related to the first two moments as

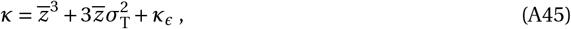

where *κ*_*ϵ*_ captures an excess of skewness compared to a normal distribution.

Using the closure of eq. (A45) in eq. (A44) and the fact that 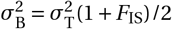, we have

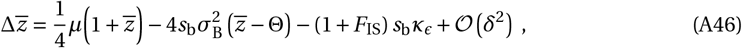

where we identify three terms capturing: (i) the effects of saltatory amplification (similar to eq. 13a); (ii) the effects of directional selection due to stabilizing selection towards Θ (increasing 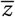 when 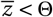, and decreasing 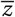 when 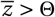); and lastly (iii) the interaction of skewness with stabilizing selection: when the distribution has a positive excess skewness (*κ*_*ϵ*_ > 0; in other words, the distribution is long-tailed), stabilizing selection is more efficient purging larger-than-average sequences than shorter-than-average ones, causing an reduction in mean (and conversely when *κ*_*ϵ*_ *<* 0).

Finally, we use the stationary variance as a function of 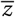 (eq. 14, calculated under neutrality) into eq. (A46) and solve it for 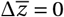. For a general excess skewness *κ*_*ϵ*_ > 0, the expression for 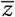 is large and complicated and we therefore refrain from showing it here. However, for a normal closure (*κ*_*ϵ*_ *=* 0), we obtain 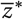 as presented in eq. (20) of the main text.

### B. Computer simulations on SLiM

The SLiM (v. 4.3; Haller and Messer, 2023) code used for our simulations is available at: https://vsudbrack.github.io/TandemRepeatsSM

These simulations track the TRs length in each homologous chromosome of each individual in a population with a given selfing rate (implemented in SLiM with the `selfingRate` property of populations).

The life-cycle described in Sec. 2.1 is implemented on SLiM using custom callbacks and properties as follows. Each diploid individual carries two genomes, each containing a single chromosome position At this position, a mutation of type `m1` is present, and its `selectionCoeff` property encodes the integer length of a tandem repeat (TR): *z*_*i*1_ for `genome1` and *z*_*i*2_ for `genome2`.

To implement individual-specific reproductive success, we assign fecundity values directly by setting each individual’s `fitnessScaling` property. We suppress SLiM default fitness calculation for mutations `m1` with `mutationEffect(m1)` callback. Fecundity is then computed from the TR lengths according to one of the functions defined in the main text (e.g., eqs. 12, 17, 18, or 19).

Offspring are then generated from adults proportionally to their `fitnessScaling`, following SLiM default reproduction mechanism. Gamete production includes saltatory amplification of TRs, which is modelled via the `mutation(m1)` callback. New mutations are generated at position `0`, with the new TR length drawn from the uniform distribution described in eq. (9), and replace the previous allele (using `mutationStackPolicy` as last).

During offspring production, unequal recombination is implemented via the `modifyChild()` call-back. Parental gametes are randomly selected, and its TR length is modified to simulate unequal recombination with a given probability, and thus we produce a new allele to the offspring following the distribution of eq. (10).

The initial conditions set on generation 1 are a monomorphic population, where all TRs start with an equal initial value *z*_0_, which we take as the closest integer to

- 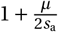 when selection is purifying (in secs. 3.1 and 3.4);
- 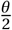 when selection has epistatic interactions;
- 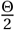 when selection is stabilizing.

We let the population evolve during a transient phase before start recording the values in the population.

Starting at generation 5^*′*^001, we measure at every 200 generations: the mean (eq. 1), the variances within and between individuals with (eq. 2), skewness (based on eq. A1 with *n =* 3), kurtosis (based on eq. A1 with *n =* 4) and genetic load (eq. 8). In selected simulations we also recorded all trait values in the population to produce histograms. We record the data until at least 50^*′*^000 generations. Each simulation had multiple independent populations replicated with the same selfing rate (4 replicates for *N =* 5^*′*^000 and 10 replicates for *N =* 2^*′*^000).

### Supplementary Figures

**Supplementary Figure S1:**
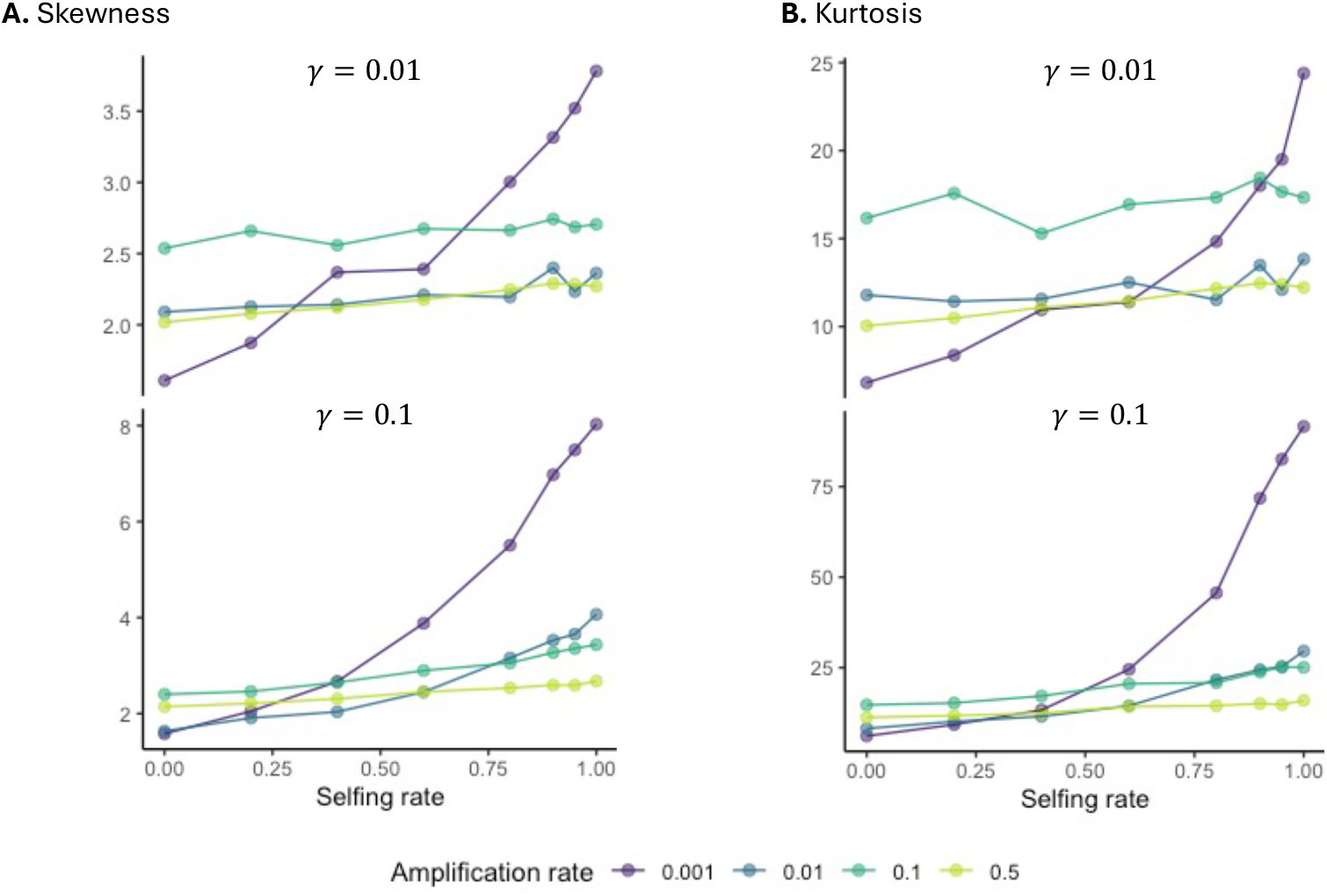
**A**. Skewness 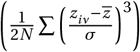 in TRs length distribution for different selfing and amplification rates (see legend). Connected points are averages of the within-population skewness across simulations after reaching equilibrium (Appendix B for details). Parameters: *N =* 2^*′*^000, *s*_a_ *=* 0.0 01 and *γ =* 0.0 1 in top row, while *γ =* 0.1 in bottom row. **B**. Similar to A., but showing kurtosis 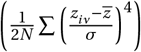. Parameters: same as A.

**Supplementary Figure S2:**
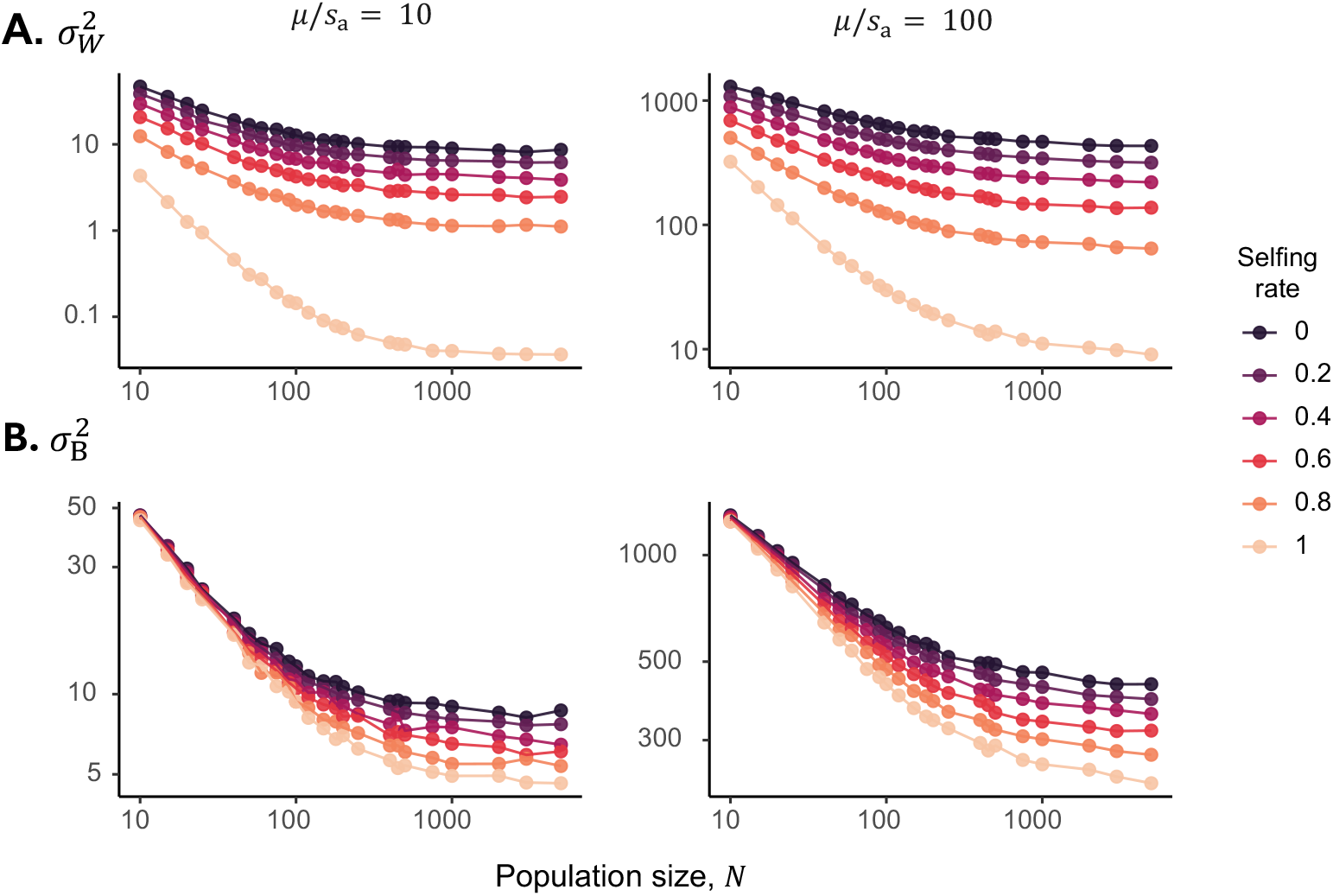
**A**. Variance within individuals 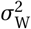 and **B**. variance between individuals 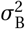 (both in logscale) within populations of size *N* for various selfing rates (see legend). The number of replicates varies between population sizes, with at least 10 simulations for each parameter combination. Parameters: same as fig. 3.

**Supplementary Figure S3:**
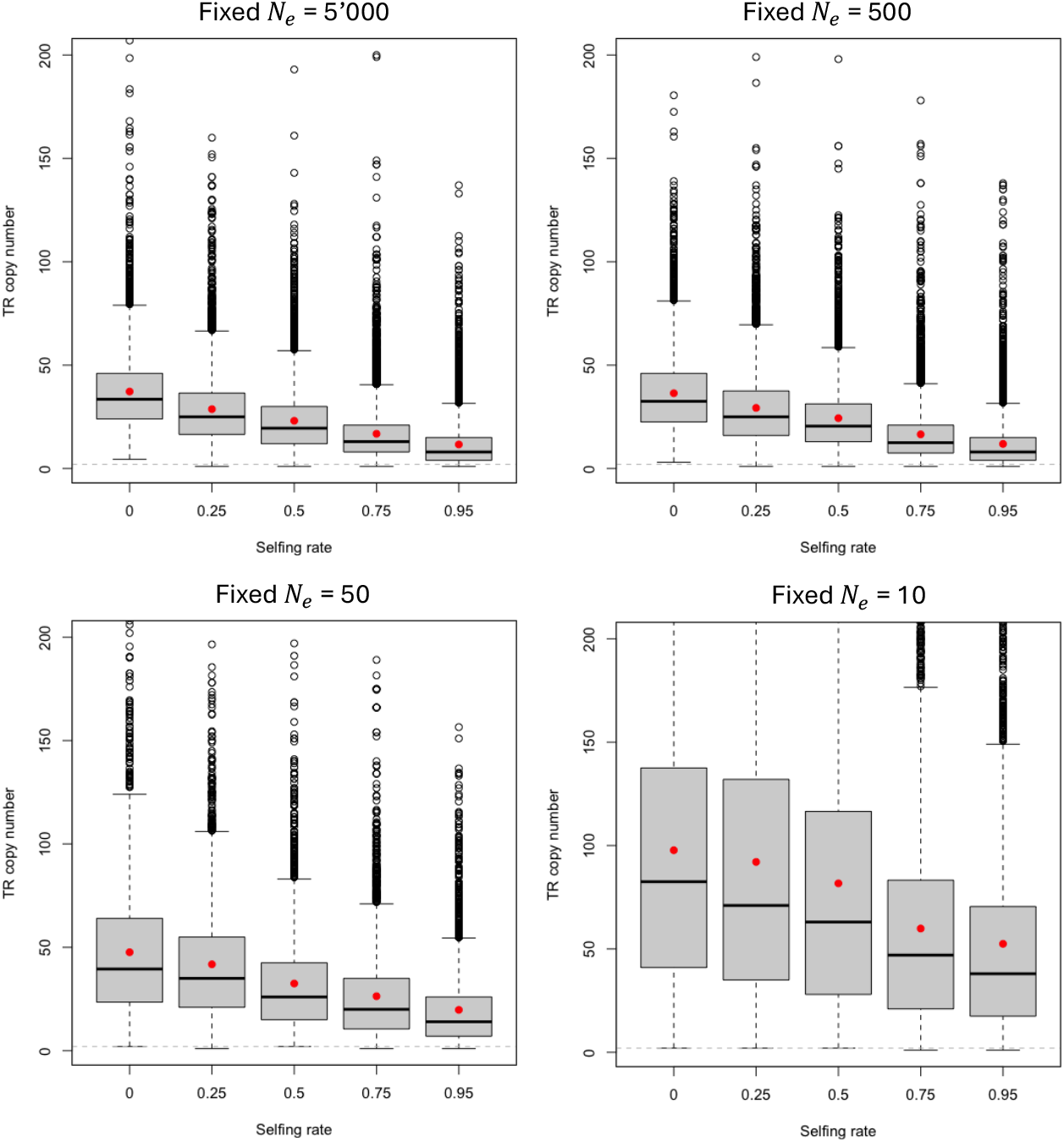
Simulations where the population size is changed such that the effective population size remains the same under various selfing rates. We set *N* = [*N*_e_ (1 *α*/2)] (eq. 6 of Pollak, 1987), where [·] means rounding to the closest integer. Each boxplot shows the median (horizontal line), interquartile range (box), and the full data spread (outliers) of TRs length from simulations. Red dots are the mean TRs length 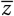 within populations. These show that even after controlling for effective population size, selfing still reduces the mean TRs length in all populations. Parameters: *γ =* 0.1, *µ =* 0.1 and *s*_a_ *=* 0.001.

**Supplementary Figure S4:**
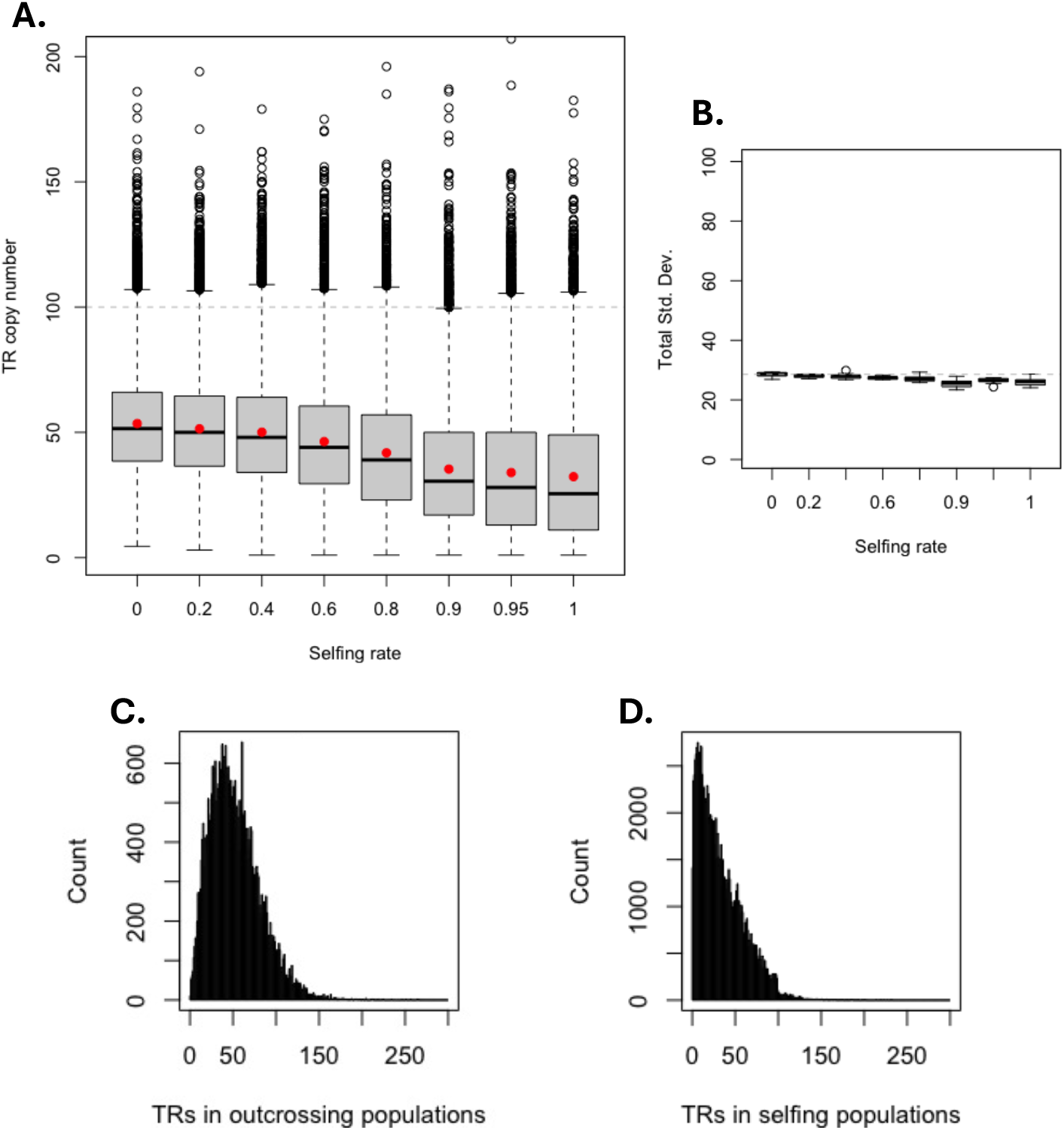
**A**. Statistics of the mean number of TRs length across different selfing rates under strongly non-additive purifying selection (close to truncation-selection, *s*_*ϵ*_ *=* 50). Boxplots display the minimum, first quartile, median, third quartile, and maximum values. The red dot is the mean across all population with given selfing rate. **B**. Total standard deviation *σ*_T_ for each selfing rate. Boxplots show values of standard deviation across the 10 replicates. We note very little variation between replicates. **C**. Histogram of TR copy number *z*_*iν*_ across populations with *α <* 0.1. **D**. Histogram of TR copy number *z*_*iν*_ across populations with *α* ≥ 0.9. Parameters: *γ =* 0.1, *µ =* 0.1, *θ =* 100 (dashed horizontal line in A) and *N =* 2^*′*^000.

**Supplementary Figure S5:**
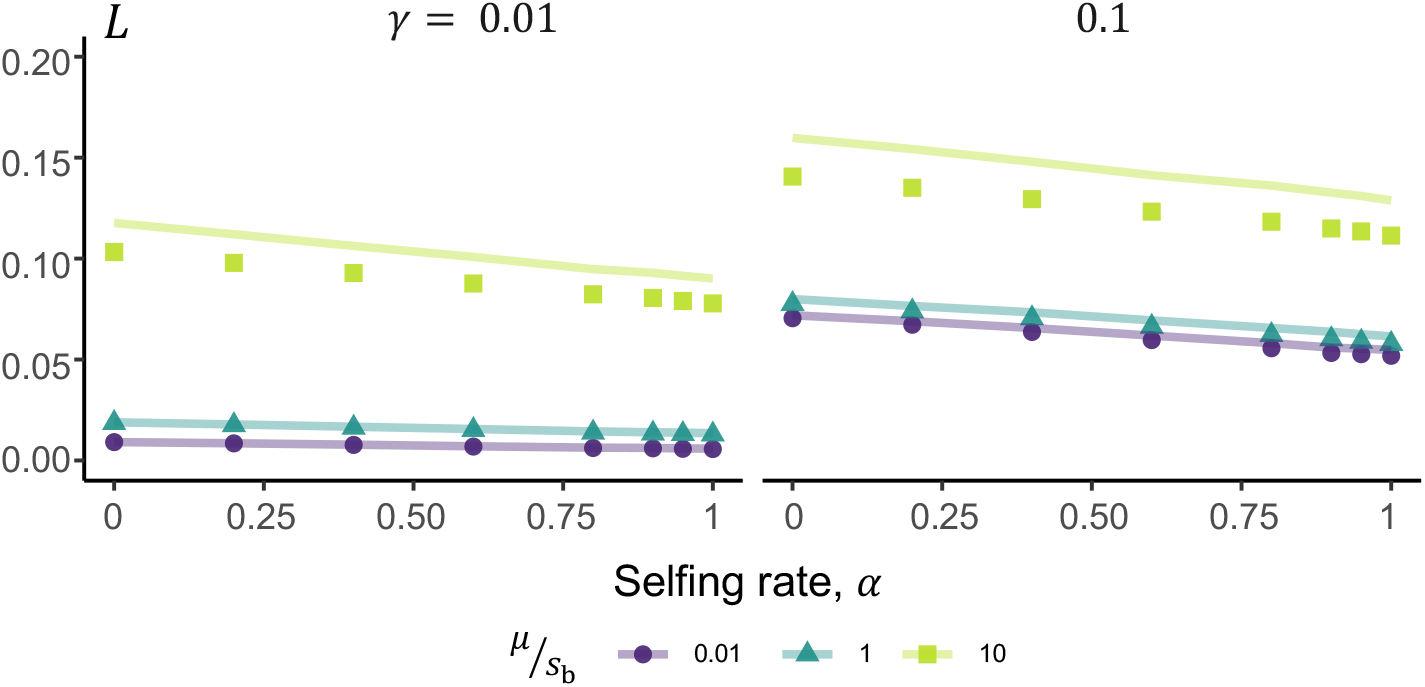
Mean load *L* due to TRs under stabilizing selection, calculated from the simulation with eq. (8), where *f*_max_ = 1. Curves represent the approximation 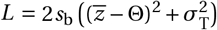, using the 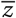 and 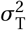 from the simulations. Parameters: *N =* 5^*′*^000, Θ *=* 10 and *s*_b_ *=* 0.01.

## References

Abu Awad, D. and Roze, D. (2018). Effects of partial selfing on the equilibrium genetic variance, mutation load, and inbreeding depression under stabilizing selection. Evolution, 72(4):751–769.

Balzano, E., Pelliccia, F., and Giunta, S. (2021). Genome (in) stability at tandem repeats. In Seminars in cell & developmental biology, volume 113, pages 97–112. Elsevier.

Burgarella, C. and Glémin, S. (2017). Population genetics and genome evolution of selfing species. eLS.

Buschiazzo, E. and Gemmell, N. J. (2006). The rise, fall and renaissance of microsatellites in eukaryotic genomes. Bioessays, 28(10):1040–1050.

Caballero, A. and Hill, W. G. (1992). Effects of partial inbreeding on fixation rates and variation of mutant genes. Genetics, 131(2):493–507.

Charlesworth, B., Sniegowski, P., and Stephan, W. (1994). The evolutionary dynamics of repetitive dna in eukaryotes. Nature, 371(6494):215–220.

Cooke, H. (1976). Repeated sequence specific to human males. Nature, 262(5565):182–186.

Crow, J. F. and Kimura, M. (1970). An introduction to population genetics theory. Scientific Publishers.

Depienne, C. and Mandel, J.-L. (2021). 30 years of repeat expansion disorders: What have we learned and what are the remaining challenges? The American Journal of Human Genetics, 108(5):764–785.

Duitama, J., Zablotskaya, A., Gemayel, R., Jansen, A., Belet, S., Vermeesch, J. R., Verstrepen, K. J., and Froyen, G. (2014). Large-scale analysis of tandem repeat variability in the human genome. Nucleic acids research, 42(9):5728–5741.

Ellegren, H. (2002). Microsatellite evolution: a battle between replication slippage and point mutation. TRENDS in Genetics, 18(2):70.

Fan, H. and Chu, J.-Y. (2007). A brief review of short tandem repeat mutation. Genomics, proteomics & bioinformatics, 5(1):7–14.

Fotsing, S. F., Margoliash, J., Wang, C., Saini, S., Yanicky, R., Shleizer-Burko, S., Goren, A., and Gymrek, M. (2019). The impact of short tandem repeat variation on gene expression. Nature genetics, 51(11):1652–1659.

Glémin, S. (2007). Mating systems and the efficacy of selection at the molecular level. Genetics, 177(2):905–916.

Goldstein, D. B., Ruiz Linares, A., Cavalli-Sforza, L. L., and Feldman, M. W. (1995). An evaluation of genetic distances for use with microsatellite loci. Genetics, 139(1):463–471.

Gymrek, M., Willems, T., Guilmatre, A., Zeng, H., Markus, B., Georgiev, S., Daly, M. J., Price, A. L., Pritchard, J. K., Sharp, A. J., et al. (2016). Abundant contribution of short tandem repeats to gene expression variation in humans. Nature genetics, 48(1):22–29.

Haller, B. C. and Messer, P. W. (2023). Slim 4: multispecies eco-evolutionary modeling. The American Naturalist, 201(5):E127–E139.

Hammond, H. A., Jin, L., Zhong, Y., Caskey, C. T., and Chakraborty, R. (1994). Evaluation of 13 short tandem repeat loci for use in personal identification applications. American journal of human genetics, 55(1):175.

Hartfield, M. and Glémin, S. (2014). Hitchhiking of deleterious alleles and the cost of adaptation in partially selfing species. Genetics, 196(1):281–293.

Jeffreys, A. J., Wilson, V., and Thein, S. L. (1985). Hypervariable ‘minisatellite’regions in human dna. Nature, 314(6006):67–73.

John, B. and Miklos, G. L. G. (1979). Functional aspects of satellite dna and heterochromatin. International review of cytology, 58:1–114.

Khristich, A. N. and Mirkin, S. M. (2020). On the wrong dna track: Molecular mechanisms of repeatmediated genome instability. Journal of Biological Chemistry, 295(13):4134–4170.

Knox, M. A., Biggs, P. J., Garcia-R, J. C., and Hayman, D. T. (2024). Quantifying replication slippage error in cryptosporidium metabarcoding studies. The Journal of Infectious Diseases, page jiae065.

Krüger, J. and Vogel, F. (1975). Population genetics of unequal crossing over. Journal of Molecular Evolution, 4:201–247.

Kruglyak, S., Durrett, R., Schug, M. D., and Aquadro, C. F. (2000). Distribution and abundance of microsatellites in the yeast genome can be explained by a balance between slippage events and point mutations. Molecular Biology and Evolution, 17(8):1210–1219.

Kruglyak, S., Durrett, R. T., Schug, M. D., and Aquadro, C. F. (1998). Equilibrium distributions of microsatellite repeat length resulting from a balance between slippage events and point mutations. Proceedings of the National Academy of Sciences, 95(18):10774–10778.

Kuderna, L. F., Gao, H., Janiak, M. C., Kuhlwilm, M., Orkin, J. D., Bataillon, T., Manu, S., Valenzuela, A., Bergman, J., Rousselle, M., et al. (2023). A global catalog of whole-genome diversity from 233 primate species. Science, 380(6648):906–913.

Lareu, M., Barral, S., Salas, A., Pestoni, C., and Carracedo, A. (1998). Sequence variation of a hypervariable short tandem repeat at the d1s1656 locus. International journal of legal medicine, 111(5):244–247.

Legendre, M., Pochet, N., Pak, T., and Verstrepen, K. J. (2007). Sequence-based estimation of minisatellite and microsatellite repeat variability. Genome research, 17(12):1787–1796.

Levinson, G. and Gutman, G. A. (1987). Slipped-strand mispairing: a major mechanism for dna sequence evolution. Molecular biology and evolution, 4(3):203–221.

Lundström, O. S., Verbiest, M. A., Xia, F., Jam, H. Z., Zlobec, I., Anisimova, M., and Gymrek, M. (2023). Webstr: a population-wide database of short tandem repeat variation in humans. Journal of molecular biology, 435(20):168260.

McGinty, R. J., Balick, D. J., Mirkin, S. M., and Sunyaev, S. R. (2025). Inherent instability of simple dna repeats shapes an evolutionarily stable distribution of repeat lengths. bioRxiv.

Miklos, G. L. G. and Gill, A. C. (1982). Nucleotide sequences of highly repeated dnas; compilation and comments. Genetics Research, 39(1):1–30.

Ohta, T. (1981). Population genetics of selfish dna. Nature, 292(5824):648–649.

Ohta, T. (1983a). On the evolution of multigene families. Theoretical population biology, 23(2):216– 240.

Ohta, T. (1983b). Theoretical study on the accumulation of selfish dna. Genetics Research, 41(1):1–15.

Ohta, T. and Kimura, M. (1981). Some calculations on the amount of selfish dna. Proceedings of the National Academy of Sciences, 78(2):1129–1132.

Perelson, A. S. and Bell, G. I. (1977). Mathematical models for the evolution of multigene families by unequal crossing over. Nature, 265(5592):304–310.

Pollak, E. (1987). On the theory of partially inbreeding finite populations. i. partial selfing. Genetics, 117(2):353–360.

Prado-Martinez, J., Sudmant, P. H., Kidd, J. M., Li, H., Kelley, J. L., Lorente-Galdos, B., Veeramah, K. R., Woerner, A. E., O’connor, T. D., Santpere, G., et al. (2013). Great ape genetic diversity and population history. Nature, 499(7459):471–475.

Pumpernik, D., Oblak, B., and Borštnik, B. (2008). Replication slippage versus point mutation rates in short tandem repeats of the human genome. Molecular Genetics and Genomics, 279:53–61.

Quilez, J., Guilmatre, A., Garg, P., Highnam, G., Gymrek, M., Erlich, Y., Joshi, R. S., Mittelman, D., and Sharp, A. J. (2016). Polymorphic tandem repeats within gene promoters act as modifiers of gene expression and dna methylation in humans. Nucleic acids research, 44(8):3750–3762.

Richard, G.-F., Kerrest, A., and Dujon, B. (2008). Comparative genomics and molecular dynamics of dna repeats in eukaryotes. Microbiology and molecular biology reviews, 72(4):686–727.

Rousset, F. (2004). Genetic structure and selection in subdivided populations, volume 40. Princeton University Press.

Rubinsztein, D. C., Amos, W., Leggo, J., Goodburn, S., Jain, S., Li, S.-H., Margolis, R. L., Ross, C. A., and Ferguson-Smith, M. A. (1995). Microsatellite evolution—evidence for directionality and variation in rate between species. Nature genetics, 10(3):337–343.

Sianta, S. A., Peischl, S., Moeller, D. A., and Brandvain, Y. (2023). The efficacy of selection may increase or decrease with selfing depending upon the recombination environment. Evolution, 77(2):394– 408.

Slatkin, M. (1995). A measure of population subdivision based on microsatellite allele frequencies. Genetics, 139(1):457–462.

Smith, G. P. (1976). Evolution of repeated dna sequences by unequal crossover: Dna whose sequence is not maintained by selection will develop periodicities as a result of random crossover. Science, 191(4227):528–535.

Srivastava, S., Avvaru, A. K., Sowpati, D. T., and Mishra, R. K. (2019). Patterns of microsatellite distribution across eukaryotic genomes. BMC genomics, 20:1–14.

Stephan, W. (1986). Recombination and the evolution of satellite dna. Genetics Research, 47(3):167– 174.

Stephan, W. (1987). Quantitative variation and chromosomal location of satellite dnas. Genetics Research, 50(1):41–52.

Stephan, W. (1989). Tandem-repetitive noncoding dna: forms and forces. Molecular biology and evolution, 6(2):198–212.

Stetsenko, R. and Roze, D. (2022). The evolution of recombination in self-fertilizing organisms. Genetics, 222(1):iyac114.

Takahata, N. (1981). A mathematical study on the distribution of the number of repeated genes per chromosome. Genetics Research, 38(1):97–102.

Usdin, K., House, N. C., and Freudenreich, C. H. (2015). Repeat instability during dna repair: Insights from model systems. Critical reviews in biochemistry and molecular biology, 50(2):142–167.

Verbiest, M., Maksimov, M., Jin, Y., Anisimova, M., Gymrek, M., and Bilgin Sonay, T. (2023). Mutation and selection processes regulating short tandem repeats give rise to genetic and phenotypic diversity across species. Journal of evolutionary biology, 36(2):321–336.

Viard, F., Bremond, P., Labbo, R., Justy, F., Delay, B., and Jarne, P. (1996). Microsatellites and the genetics of highly selfing populations in the freshwater snail bulinus truncatus. Genetics, 142(4):1237– 1247.

Walsh, B. and Lynch, M. (2018). Evolution and selection of quantitative traits. Oxford University Press.

Walsh, J. B. (1987). Persistence of tandem arrays: implications for satellite and simple-sequence dnas. Genetics, 115(3):553–567.

Wei, K. H.-C., Grenier, J. K., Barbash, D. A., and Clark, A. G. (2014). Correlated variation and population differentiation in satellite dna abundance among lines of drosophila melanogaster. Proceedings of the National Academy of Sciences, 111(52):18793–18798.

Wright, S. (1949). The genetical structure of populations. Annals of eugenics, 15(1):323–354.

Wright, S. I., Ness, R. W., Foxe, J. P., and Barrett, S. C. (2008). Genomic consequences of outcrossing and selfing in plants. International Journal of Plant Sciences, 169(1):105–118.

Zenke, P., Egyed, B., Zöldág, L., and Padar, Z. (2011). Population genetic study in hungarian canine populations using forensically informative str loci. Forensic Science International: Genetics, 5(1):e31–e36.

